# Functional characterization of Atlantic salmon (*Salmo salar* L.) PepT2 transporters

**DOI:** 10.1101/2022.02.11.480090

**Authors:** Francesca Vacca, Ana S. Gomes, Koji Murashita, Raffella Cinquetti, Cristina Roseti, Amilcare Barca, Ivar Rønnestad, Tiziano Verri, Elena Bossi

## Abstract

The high-affinity/low-capacity system Slc15a2 (PepT2) is responsible for the reuptake of di/tripeptides from the renal proximal tubule, but it also operates in many other tissues/organs. Information regarding PepT2 in teleost fish is limited and to date functional data are available from the zebrafish (*Danio rerio*) only. Here, we report the identification of two *slc15a2* genes in the Atlantic salmon (*Salmo salar*) genome, namely *slc15a2a* and *slc15a2b.* The two encoded PepT2 proteins share 87% identity and resemble both structurally and functionally to the canonical vertebrate PepT2 system. The mRNA tissue distribution analyses reveal a widespread distribution of *slc15a2a* transcripts, being more abundant in the brain and gills, while *slc15a2b* transcripts are mainly expressed in kidney and distal part of gastrointestinal tract. The function of the two transporters was investigated by heterologous expression in *Xenopus laevis* oocytes and two-electrode voltage-clamp recordings of transport and presteady-state currents. Both PepT2a and PepT2b in the presence of Gly-Gln elicit pH-dependent and Na^+^ independent inward currents. The biophysical and kinetic analysis of the recorded currents defined the transport properties, confirming that the two Atlantic salmon PepT2 proteins behave as high-affinity/low-capacity transporters. The recent structures and the previous kinetic schemes of rat and human PepT2 qualitatively account for the characteristics of the two Atlantic salmon proteins. This study is the first to report on the functional expression of two PepT2-type transporters that operate in the same vertebrate organism as a result of (a) gene duplication process(es).

**Key points summary:**

- Two *slc15a2*-type genes, *slc15a2a* and *slc15a2b* coding for PepT2-type peptide transporters were found in the Atlantic salmon.
- *slc15a2a* transcripts, widely distributed in the fish tissues, are abundant in brain and gills, while *slc15a2b* transcripts are mainly expressed in kidney and distal gastrointestinal tract.
- Amino acids involved in vertebrate Slc15 transport function are conserved in PepT2a and PepT2b proteins.
- Detailed kinetic analysis indicates that both PepT2a and PepT2b operate as high-affinity transporters.
- The kinetic schemes and structures proposed for the mammalian models of PepT2 are suitable to explain the function of the two Atlantic salmon transporters.

## Introduction

The Solute carrier 15 (Slc15) family includes H^+^-dependent transporters traditionally divided into Peptide Transporters and Peptide/Histidine Transporters, all known for their key role in the cellular uptake/reuptake of di/tripeptides and peptidomimetics (Smith *et al*., 2013; Zhao & Lu, 2015; Viennois *et al*., 2018). The Peptide Transporters sub-group includes two transport systems: the low-affinity/high-capacity system Slc15a1 (PepT1), mainly expressed in the brush border membrane (BBM) of the small intestinal enterocytes, and the high-affinity/low-capacity system Slc15a2 (PepT2). PepT2 was initially discovered as the peptide transporter of the BBM of the renal proximal tubular cell but has been shown to be widely distributed in (epithelial and non-epithelial cells of) many other organs/tissues including the central nervous system, the choroid plexus, the enteric nervous system, lung, skin, intestine, glands, testis, prostate, ovary, uterus, and eye (Ruhl *et al*., 2005; Biegel *et al*., 2006). First cloned from human kidney in 1995 (Liu *et al*., 1995), PepT2 was later well described in several mammalian species. In terms of function, detailed electrophysiological data for the transporter were first reported for rabbit (Boll *et al*., 1996), and then for rat (Wang *et al*., 1998), human (Chen *et al*., 1999; Sala-Rabanal *et al*., 2008) and mouse (Rubio-Aliaga *et al*., 2000). In parallel, a very large number of di/tripeptides, peptidomimetics and peptide-like drugs have been tested and shown to be substrates of the mammalian PepT2 transporters [reviewed in e.g. (Smith *et al*., 2013); (Zhao & Lu, 2015); (Viennois *et al*., 2018)].

Despite the increased interest in piscine di/tripeptide transporters over the last years, mainly due to their role in digestive and absorptive physiology, growth and nutrition [see e.g. (Verri *et al*., 2010; Verri *et al*., 2011; Romano *et al*., 2014; Verri *et al*., 2017)], functional data on PepT2 are still missing, except for the zebrafish (*Danio rerio*) (Romano *et al*., 2006). Based on the zebrafish data, piscine PepT2 operates as a ‘canonical’ low-capacity/high-affinity transporter and is mainly expressed in the kidney, brain and intestine, but it is also present in the gills, eye, and skeletal muscle (Romano *et al*., 2006). Later studies in other teleost fish species, mostly of commercial interest, have shown that PepT2 is present in the intestine, specifically confined to the mid-to-distal part and mainly located at the BBM of the enterocytes [see e.g. (Con *et al*., 2017)]. Probably, this is to support the absorption of (residual) luminal di/tripeptides that reach the terminal gut region (summarized in **Table 1**).

**Table 1.**
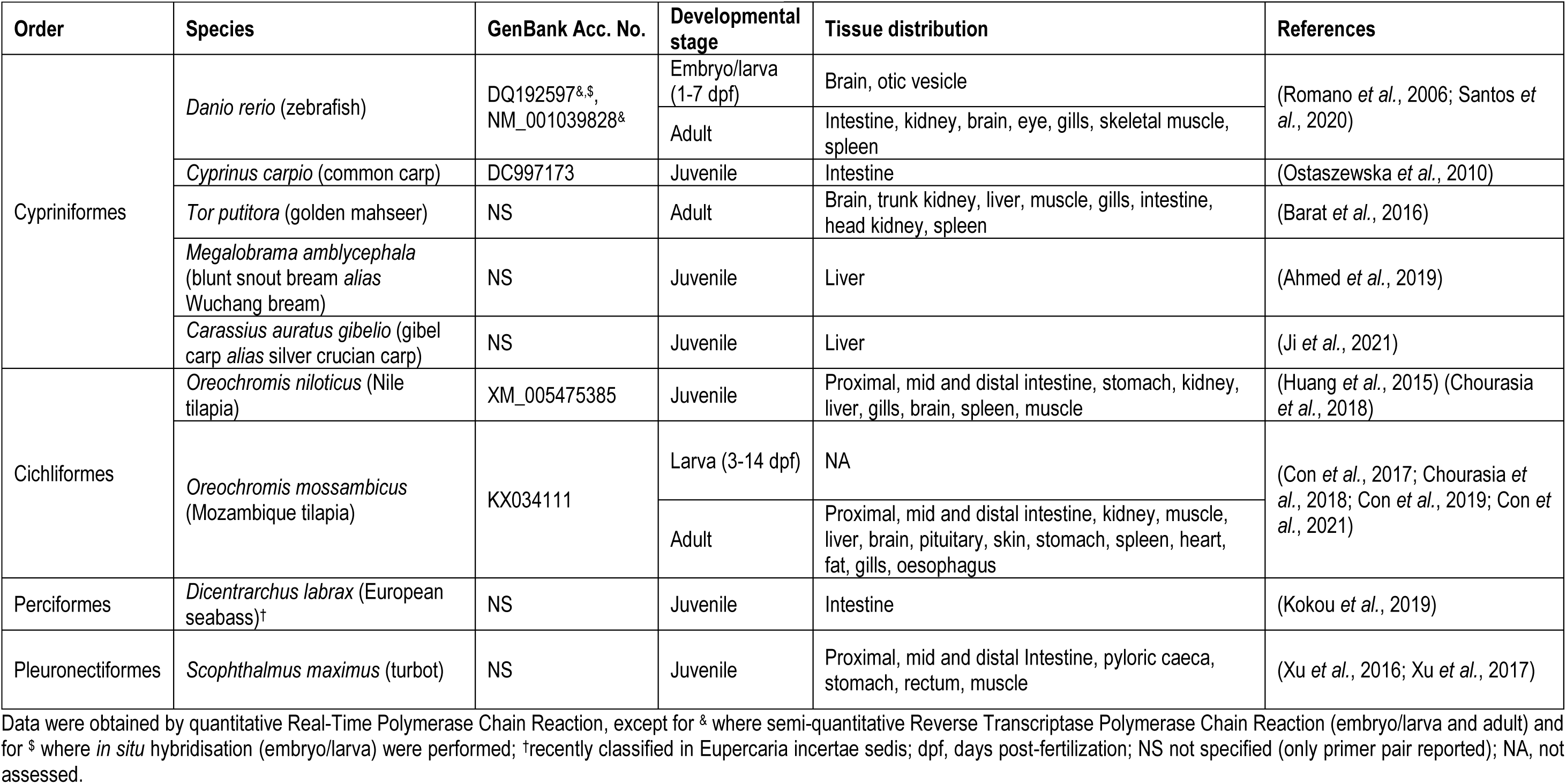
Organ/tissue distribution of *slc15a2* mRNA in teleost fish.

| Order | Species | GenBank Acc. No. | Developmental stage | Tissue distribution | References |
| --- | --- | --- | --- | --- | --- |
| Cypriniformes | *Danio rerio* (zebrafish) | DQ192597<sup>&,\$</sup>,<br>NM_001039828<sup>&</sup> | Embryo/larva (1-7 dpf) | Brain, otic vesicle | (Romano *et al.*, 2006; Santos *et al.*, 2020) |
|  |  |  | Adult | Intestine, kidney, brain, eye, gills, skeletal muscle, spleen |  |
|  | *Cyprinus carpio* (common carp) | DC997173 | Juvenile | Intestine | (Ostaszewska *et al.*, 2010) |
|  | *Tor putitora* (golden mahseer) | NS | Adult | Brain, trunk kidney, liver, muscle, gills, intestine, head kidney, spleen | (Barat *et al.*, 2016) |
|  | *Megalobrama amblycephala* (blunt snout bream *alias* Wuchang bream) | NS | Juvenile | Liver | (Ahmed *et al.*, 2019) |
|  | *Carassius auratus gibelio* (gibel carp *alias* silver crucian carp) | NS | Juvenile | Liver | (Ji *et al.*, 2021) |
| Cichliformes | *Oreochromis niloticus* (Nile tilapia) | XM_005475385 | Juvenile | Proximal, mid and distal intestine, stomach, kidney, liver, gills, brain, spleen, muscle | (Huang *et al.*, 2015) (Chourasia *et al.*, 2018) |
|  | *Oreochromis mossambicus* (Mozambique tilapia) | KX034111 | Larva (3-14 dpf) | NA | (Con *et al.*, 2017; Chourasia *et al.*, 2018; Con *et al.*, 2019; Con *et al.*, 2021) |
|  |  |  | Adult | Proximal, mid and distal intestine, kidney, muscle, liver, brain, pituitary, skin, stomach, spleen, heart, fat, gills, oesophagus |  |
| Perciformes | *Dicentrarchus labrax* (European seabass)<sup>†</sup> | NS | Juvenile | Intestine | (Kokou *et al.*, 2019) |
| Pleuronectiformes | *Scophthalmus maximus* (turbot) | NS | Juvenile | Proximal, mid and distal Intestine, pyloric caeca, stomach, rectum, muscle | (Xu *et al.*, 2016; Xu *et al.*, 2017) |

Data were obtained by quantitative Real-Time Polymerase Chain Reaction, except for <sup>&</sup> where semi-quantitative Reverse Transcriptase Polymerase Chain Reaction (embryo/larva and adult) and for <sup>\$</sup> where *in situ* hybridisation (embryo/larva) were performed; <sup>†</sup>recently classified in Eupercaria incertae sedis; dpf, days post-fertilization; NS not specified (only primer pair reported); NA, not assessed.

In this paper, we report the characterization of two newly identified PepT2-type proteins in the Atlantic salmon (*Salmo salar*, L.). Both transporters structurally and functionally resemble the mammalian PepT2 paradigm, thus differing from the two PepT1-type proteins previously described in salmon (Ronnestad *et al*., 2010; Gomes *et al*., 2020). The kinetic properties of the two Atlantic salmon PepT2 transporters have been explored by expressing the proteins in *Xenopus laevis* oocytes and then analyzed by two-electrode voltage-clamp recordings of the transport current (*I*t) and the presteady-state current (*I*_PSS_). The presence of their mRNAs has also been analyzed in several target tissues. To our knowledge, this is the first report that focuses on two *slc15a2* (paralogue) genes in a vertebrate species, a result of the specific salmonid whole-genome duplication (WGD) event, and that comparatively accesses the kinetics of two highly similar PepT2 transporters belonging to the same genetic background.

## Materials and Methods

### Ethics statement

The research using Atlantic salmon was conducted in accordance with the Norwegian Animal Welfare Act of 12 December 1974, no. 73, § 22 and § 30, amended 19 June 2009, and all handling and procedures related to fish described in this study have been approved by the National Animal Research Authority in Norway (Fots ID 14984). The Cargill Innovation Center (Dirdal, Norway) facility has a general permission to conduct fish experiments, license number 2016/2835 (24 February 2016) provided by the Norwegian Food Safety Authority.

The research involving *Xenopus laevis* oocytes was conducted using an experimental protocol approved locally by the Committee of the *“Organismo Preposto al Benessere degli Animali”* of the University of Insubria (OPBA permit no. 02_15) and by the Italian Ministry of Health (permit no. 1011/2015).

Investigators understand the ethical principles under which the journal operates and declare that their work complies with the journal animal ethics checklist.

### Animals and tissue collection

Atlantic salmon were reared (Cargill Innovation Center, Dirdal, Norway) following standard procedures, in seawater at 8.7 °C with a stock density below 25 kg fish/m^3^ and constant day light. The fish were fed a commercial EWOS 10 mm pellet size diet [for information regarding diet composition please refer to (Gomes *et al*., 2020)] using automatic feeders 3 times a day (22:00-24:00, 02:00-04:00, 06:00-08:00). Atlantic salmon (906 ± 122 g wet weight; 38.8 ± 1.8 cm total length; n = 8) were collected 2 hours after the 06:00-08:00 meal and euthanized with an overdose of MS222 (tricaine methansulfonate 300 mg/L; Norsk Medisinaldepot AS, Bergen, Norway) on site. The selected tissues were rapidly collected, snap-frozen in liquid nitrogen and stored at -80 °C until further analyses.

### Molecular cloning

Total RNA was extracted from the hindgut and whole brain of Atlantic salmon using TRI Reagent (Millipore Sigma, St. Louis, MO, USA) following the manufacturer’s instructions. To avoid contamination with genomic DNA, total RNA was treated with Turbo DNA-Free kit (Thermo Fisher Scientific, Waltham, MA, USA) according to the manufacturer’s protocol, and 1.5 μg of DNase treated RNA was used to synthesize cDNA using SuperScript III First-Strand Synthesis system for RT-PCR kit (Thermo Fisher Scientific) with Oligo (dT)20 primers according to the manufacturer’s protocol.

Atlantic salmon *slc15a2a* and *slc15a2b* transcripts were identified using the zebrafish PepT2 nucleotide sequence (GenBank Acc. No. NM_001039828.1) as a query against the Atlantic salmon genome database available in GenBank, and specific primers were designed (**Table 2**, searches in the database performed in 2019). The complete coding sequences (CDS) of *slc15a2a* and *slc15a2b* were amplified using Q5 High-Fidelity DNA polymerase (New England Biolabs, Ipswich, MA, USA) according to the manufacturer’s protocol. The following thermal program: 98 °C for 30 s; 35 cycles of 98 °C for 10 s, 64 °C for 20 s, 72 °C for 2 min; and a final step at 72 °C for 2 min was used in a GeneAmp PCR system 2700 thermal cycler (Applied Biosystems, Foster City, CA, USA). The PCR products were resolved on 1% (w/v) agarose gel and purified using E.Z.N.A. Gel Extraction Kit (Omega bio-tek, Inc, Norcross, GA, USA). *slc15a2b* was cloned into a StrataClone blunt PCR cloning vector pSC-B (Agilent Technologies, Palo Alto, CA, USA), while an additional step which added 3’ A-overhangs to the *slc15a2a* purified PCR product was performed using Taq DNA polymerase (New England Biolabs) before cloning into pCR4-TOPO vector (Thermo Fisher Scientific). Sequencing was performed at the University of Bergen Sequencing Facility (Bergen, Norway) and sequence identity confirmed by tBLASTx analysis against the Atlantic salmon genome database available in GenBank. Furtherly, Atlantic salmon PepT2 amino acid sequences were deduced using the Expasy translate tool (https://web.expasy.org/translate/). Putative transmembrane domains were drawn using the rat PepT2 annotation data at the UniProt database (UniProtKB Acc. No. Q63424). Potential *N*-glycosylation sites were predicted using NetNGlyc 1.0 server (http://www.cbs.dtu.dk/services/NetNGlyc/), and the PTR2 family proton/oligopeptide symporter signature was defined using the ScanProsite tool (de Castro *et al*., 2006).

**Table 2.**
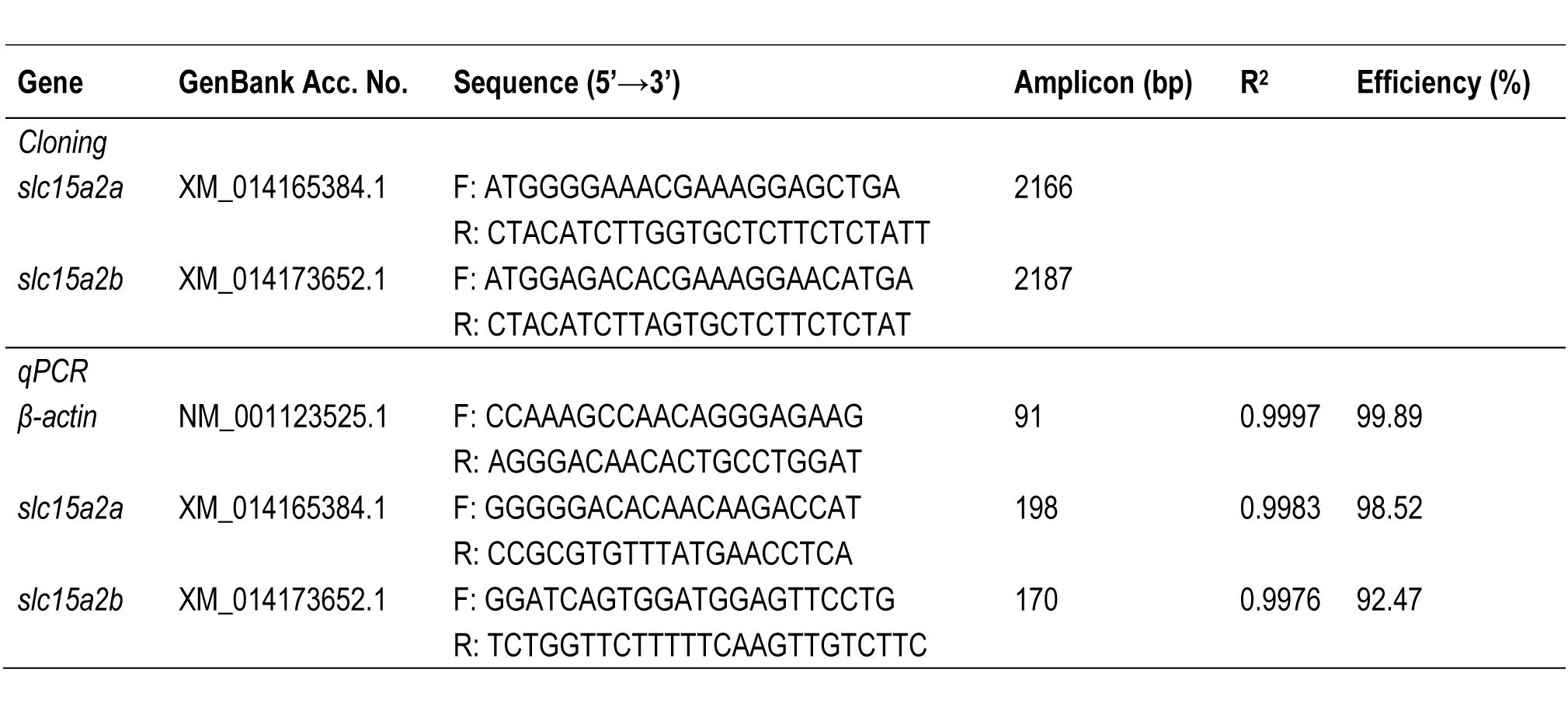
Sequence of the specific primers used for cloning and quantitative RT-PCR (qPCR) mRNA expression analysis. Sequence accession number, primer sequences, amplicon sizes, R^2^ and qPCR efficiency are indicated for each primer pair.

### Quantitative Real-Time PCR (qRT-PCR)

Total RNA was isolated from brain, gills, olfactory cavity, lip, tongue, several sections of the gastrointestinal tract (oesophagus, anterior and posterior stomach, pyloric caeca, anterior midgut, midgut, posterior midgut, and anterior and posterior hindgut), kidney and head-kidney as described above. Total RNA was treated with TURBO DNA-free (Thermo Fisher Scientific) as described above and RNA integrity assessed in all samples using an Agilent 2100 Bioanalyzer (Agilent Technologies). cDNA was synthesized as described in the section above.

Specific primers for the target transcripts and the reference gene β-actin (GenBank Acc. No. NM_001123525.1) were designed using primer-BLAST (Ye *et al*., 2012) (**Table 2**). qPCR reactions were carried out in duplicate with a final reaction volume of 20 μl using iTaq Universal SYBR Green Supermix (Bio-Rad, Hercules, CA, USA). The following qPCR conditions: 95 °C for 30 s; 40 cycles of 95 °C for 5 s, 60 °C for 25 s, were used in a CFX 96TM Real Time System (Bio-Rad). Melting curve analysis over a range of 65 to 95 °C (increment of 0.5 °C for 2 s) allowed detection of non-specific products and/or primer dimers. The assay efficiency was determined using a 10-fold dilution curve of the target gene cloned in the pCR4-TOPO vector (Thermo Fisher Scientific). The qPCR efficiency varied between ∼92% and ∼100% with a R^2^ > 0.99 (**Table 2**).

The target transcripts and β-actin copy number were calculated for each sample based on the respective standard curve, using the following expression: Copy number = (Ct - intercept) x (slope)^-1^. The target transcripts copy number was divided by the ng of total RNA used in the reaction and subsequently normalized with the β-actin copy number/ng of total RNA. Data were analyzed and figures prepared in R version 4.0.2 (Team, 2009) using the Tidyverse package (Wickham *et al*., 2019).

### Phylogeny and synteny analysis

The deduced mature protein sequences of Atlantic salmon PepT2a (GenBank Acc. No. XP_014020859.1) and PepT2b (GenBank Acc. No. XP_014029127.1), zebrafish PepT2 (UniProtKB Acc. No. B0S6T2), rat PepT2 (UniProtKB Acc. No. Q63424) and human PepT2 (UniProtKB Acc. No. Q16348) were aligned using ClustalX 2.1 with the default parameters (Gonnet series matrix, Gap opening penalty 10, Gap extension 0.2) and the percentages of similarity/identity between sequences were calculated using GeneDoc software (https://genedoc.software.informer.com/2.7/).

For phylogenetic analysis of the PepT2 family members, alignments were performed in MEGA X (Kumar *et al*., 2018) using MUSCLE with the default parameters (UPGMA clustering method, Gap opening penalty -2.90, Gap extension 0.0). The sequence alignment was analyzed for the best-fit substitution model in MEGA X to select the best statistical model to study protein family evolution. The phylogenetic tree was constructed using Maximum Likelihood (ML) with a Jones-Taylor-Thornton (JTT) model (Jones *et al*., 1992) with fixed Gamma distribution (+G) parameter with five rate categories and 1000 bootstrap replicates. Included in the analysis were the PepT2 sequences from four salmonids [Atlantic salmon, rainbow trout (*Oncorhynchus mykiss*), chinook salmon (*Oncorhynchus tshawytscha*), Arctic char (*Salvelinus alpinus*)], two cyprinids [zebrafish, goldfish (*Carassius auratus*)], medaka (*Oryzias latipes*), Nile tilapia (*Oreochromis niloticus*), turbot, Atlantic herring (*Clupea harengus*), cavefish (*Astyanax mexicanus*), the primitive freshwater ray-finned fish spotted gar (*Lepisosteus oculatus*) and human (*Homo sapiens*).

The gene environment of Atlantic salmon *slc15a2* genes was characterized and compared to the homologue *slc15a2* genome region in Northern pike and zebrafish. The genes flanking *slc15a2* were identified using the annotation provided by direct Genomicus genome browser consulting (https://www.genomicus.biologie.ens.fr/genomicus).

### Protein modeling

SWISS-MODEL (Waterhouse *et al*.) (https://swissmodel.expasy.org/interactive) was used to predict the structure of the Atlantic salmon PepT2a and PepT2b proteins using their primary sequences (GenBank Acc. Nos.: XP_014020859.1 and XP_014029127.1, respectively). The rat PepT2 (Protein Data Bank Acc. No.: 7nqk.1) (Parker *et al*., 2021) and human PepT2 (Protein Data Bank Acc. No.: 7pmy.1) (Killer *et al*., 2021) were used as templates to build the models. The final three-dimensional structures were visualized using SWISS-MODEL view tools.

### Expression in Xenopus laevis oocytes and electrophysiology

The complete open reading frame encoding for Atlantic salmon PepT2a and PepT2b was subcloned into pSPORT1 plasmid. Sequence analysis confirmed a 99% identity of PepT2a to the public available sequence (GenBank Acc. No. XM_014165384.1; two nucleotides change, and one of these alters Leu to Ser in position 9; see **Fig. 1*A***) and a 100% identity of PepT2b to the published sequence (GenBank Acc. No. XM_014173652.1).

**Figure 1.**
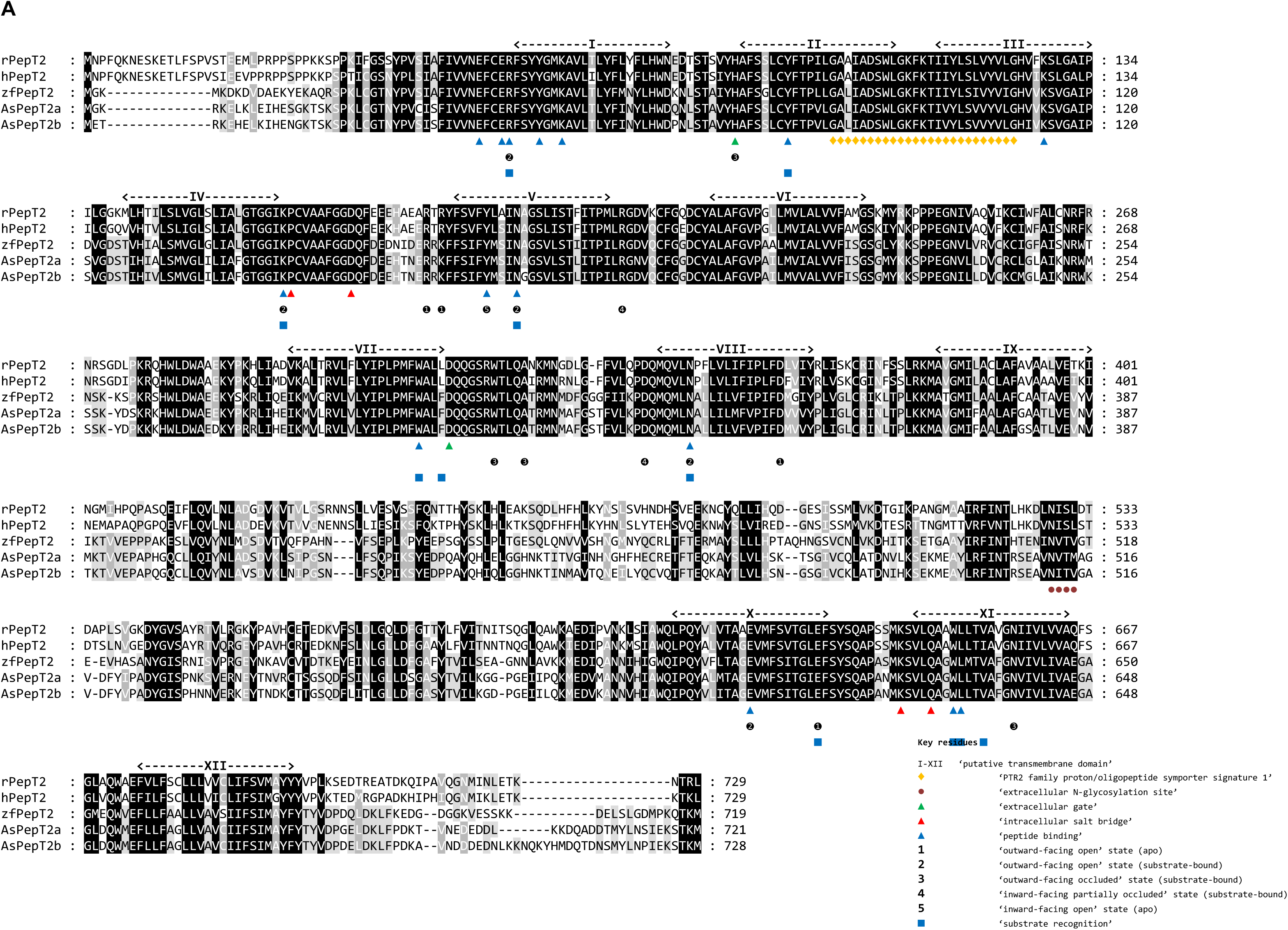
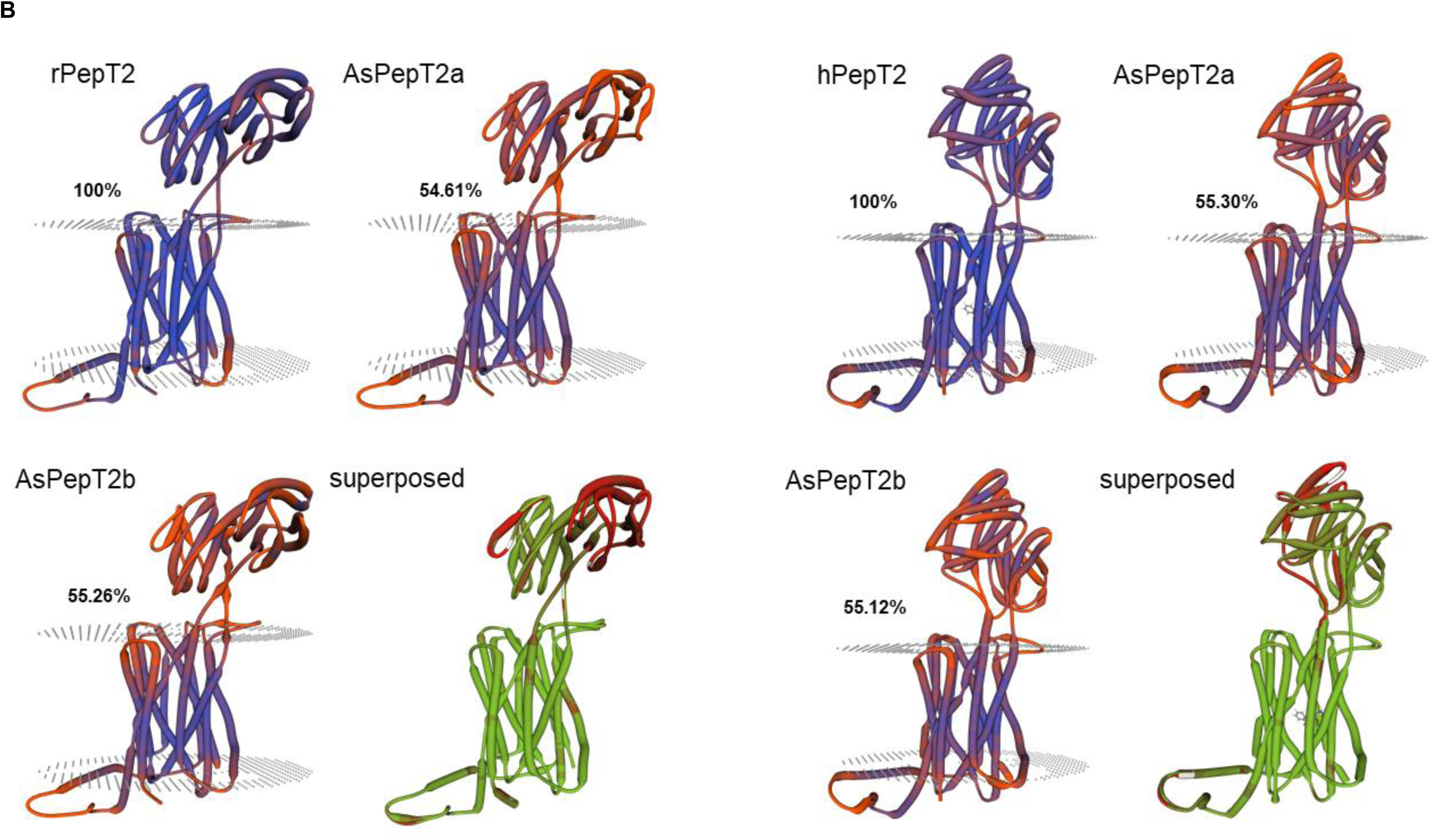
(***A***) Multiple alignments of the Atlantic salmon PepT2a (AsPepT2a) and PepT2b (AsPepT2b), zebrafish PepT2 (zfPepT2), rat PepT2 (rPepT2) and human (hPepT2) amino acid sequences as obtained by using ClustalX 2.1 and edited in GeneDoc 2.7 software. The conserved PTR2 family proton/oligopeptide symporter signature 1 (PROSITE pattern PS01022 - amino acid residues 100-124 on rPepT2 and hPepT2, and 86-110 on zPepT2, AsPepT2a and AsPepT2b) is marked by orange diamonds (♦). The putative transmembrane domains, named I to XII, were drawn using the annotation data of rPepT2 (see UniProtKB Acc. No. Q63424). Only one conserved extracellular N-glycosylation site (PROSITE pattern PS00001 – amino acid residues 528-532 on rPepT2 and hPepT2, 513-516 on zPepT2, and 511-514 on AsPepT2a and AsPepT2b), as obtained using NetNGlyc 1.0 server, is reported and marked by brown circles (●). Highlighted are key residues referring to the ‘extracellular gate’ (green triangles, ▴), ‘intracellular salt bridge’ (red triangles, ▴), and ‘peptide binding’ (blue triangles, ▴), as defined on the Cryo-EM structure of the rat PepT2 transporter (Protein Data Bank Acc. No. 7nqk.1, (Parker *et al*., 2021)). Key residues referring to the mechanism for substrate recognition and transport, as defined on the Cryo-EM structure of the human PepT2 (Protein Data Bank Acc. No. 7pmy.1) or human PepT1 (Protein Data Bank Acc. Nos. 7pn1.1, 7pmx.1 and 7pmw.1, (Killer *et al*., 2021)) are also highlighted. Amino acid residues involved in the subsequent steps of the transport cycle are shown: ‘outward-facing open’ state (apo) (**1**) (adapted from the human PepT1 structure, ref. Protein Data Bank Acc. No. 7pn1.1), ‘outward-facing open’ state (substrate-bound) (**2**) (adapted from the human PepT1 structure, ref. Protein Data Bank Acc. No. 7pmx1.1), ‘outward-facing occluded’ state (substrate-bound) (**3**) (adapted from the human PepT1 structure, ref. Protein Data Bank Acc. No. 7pmw1.1), ‘inward-facing partially occluded’ state (substrate-bound) (**4**) (adapted from the human PepT2 structure, ref. Protein Data Bank Acc. No. 7pmy1.1), ‘inward-facing open’ state (apo) (**5**) (from the human PepT1 Alphafold structure prediction, ref. Alphafold Acc. No. AF- P46059-F1;(Jumper *et al*., 2021)). The amino acid residues involved in ‘substrate recognition’ are all specifically indicated, marked by blue squares (▪) (from the human PepT2 structure, ref. Protein Data Bank Acc. No. 7pmy1.1). Substrate: Ala-Phe. For details on transport dynamics, mechanism, states and conformations, please see (Killer *et al*., 2021). (***B***) Three-dimensional representation of single and superposed structures of rPepT2, AsPepT2a and AsPepT2b (‘outward-facing open’ conformation) (left), and hPepT2, AsPepT2a and AsPepT2b (‘inward-facing partially occluded’ conformation) (substrate-bound; the substrate Ala-Phe is represented) (right), as obtained by using SWISS-MODEL tools. The models were built using as a template the rPepT2 (Protein Data Bank Acc. No. 7nqk.1) or the hPepT2 (Protein Data Bank Acc. No. 7pmy.1); then, the models were superposed by using the ‘Compare’ view tool. Colour schemes for single structures based on (model) confidence (it evaluates, for each residue of the model, the expected similarity to the native structure, thus representing an index of the ‘local quality’ of the residue): red, low confidence and blue, high confidence. Colour scheme for superposed structures based on consistency (it identifies local deviations of a protein structure from the ‘consensus’ established by all other structures selected for comparison): red, low consistency and green, high consistency. Percent identities of Atlantic salmon PepT2 proteins vs. the rat (left) and human (right) PepT2 proteins are reported.

To improve the expression in the membrane of *Xenopus laevis* oocytes, a 3’ UTR sequence of 1725 bp containing two poly-adenylation signals and a poly(A) tail from rat Divalent metal transporter 1 (rDmt1, *alias* rat *Slc11a2*; GenBank Acc. No. NM_013173.2) (Buracco *et al*., 2015) was added to the end of each Atlantic salmon PepT2 CDS.

*Xenopus laevis* oocytes were collected under anaesthesia [MS222; 0.10 % (w/v) solution in tap water] by laparotomy from adult females (Envigo, San Pietro al Natisone, Italy) and prepared as described previously (Bossi *et al*., 2007). *Xenopus laevis* frogs were maintained according to international guidelines (Delpire *et al*., 2011; McNamara *et al*., 2018). In brief, the frogs were kept in a XenoPlus amphibian System, with a continuous recirculating water system (Tecniplast, Buguggiate, Varese, Italy), and a daylight cycle of 12 hours. They were fed *ad libitum* with a *Xenopus* diet (Mucedola, Settimo Milanese, Milan, Italy). The oocytes were collected three times from each frog, and the animals were killed with an overdose of anaesthetic (0.5% (w/v) MS222) after the third oocytes collection (Torreilles *et al*., 2009).

Capped cRNAs of the two Atlantic salmon PepT2 were synthesized by *in vitro* transcription using T7 RNA polymerase from cDNAs in pSPORT1 linearized with NotI and purified with Wizard SV Gel and PCR clean-up system (Promega Italia, Milan, Italy). The purified cRNAs were quantified by NanoDrop^TM^ 2000 Spectrophotomer (Thermo Fisher Scientific), and 12.5 ng injected into the oocytes using a manual microinjection system (Drummond Scientific Company, Broomall, PA, USA). Before electrophysiological studies, the cRNA-injected oocytes were incubated at 18 °C for 3-4 days in NDE (NaCl 96 mmol/L, KCl 2 mmol/L, CaCl2 1.8 mmol/L, MgCl2 1 mmol/L, HEPES 5 mmol/L, pyruvate 2.5 mmol/L and gentamycin sulphate 0.05 mg/mL pH 7.6).

Two-electrode voltage-clamp experiments were performed using a commercial amplifier (Oocyte Clamp OC-725B, Warner Instruments, Hamden, CT, USA) and the pCLAMP software (Version 10.7, Molecular Devices, San Jose CA, CA, USA).

The holding potential was kept at -60 mV; the voltage pulse protocol consisted of 10 square pulses from -140 to +20 mV (20 mV increment) of 400 ms each. Signals were filtered at 0.1 kHz, sampled at 0.2 kHz or 0.5 kHz, and 1 kHz. Transport-associated currents (Itr) were calculated by subtracting the traces in the absence of substrate from those in its presence. When the normalization is reported the values of the steady-state currents were divided by the value of the current recorded for each transporter at -140 mV in the presence of 1 mmol/L Gly-Gln at pH 7.6. The number of samples “n” corresponds to the number of oocytes used in each condition and the batches correspond to the animals from which the oocytes were collected. These data are summarized in figure captions.

### Data analysis and figure preparation

Steady-state transport currents from substrate dose-response experiments were fitted with the Michaelis-Menten equation [1]:

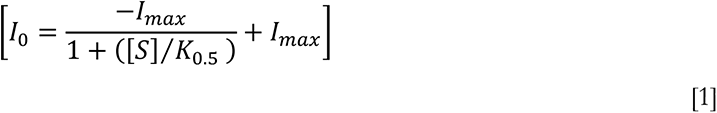

for which *I0* is the evoked current, *I*_max_ is the derived relative maximal current, S is the substrate (Gly-Gln) concentration and *K*_0.5_ is the substrate concentration at which current is half-maximal.

To analyze the pre-steady-state currents, the current traces were fitted with double-exponential function [2]:

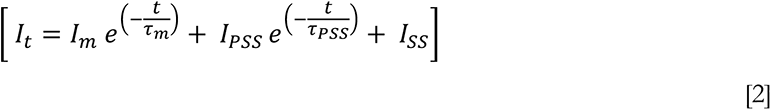

where *I*t is the total current across the oocyte membrane; *t* is the time; *I*m is a capacitive current with time constant *τ*m associated with the oocyte plasma membrane, *I*_PSS_ is a transient current associated with the transporter expression with time constant *τ*, and *I*SS is the steady-state current. At each voltage, the amount of displaced charge (*Q*) was calculated by integrating the isolated traces after zeroing any residual steady-state transport current. Finally, the charge *vs.* voltage relationship (*Q/V*) was fitted with the Boltzmann equation [3]:

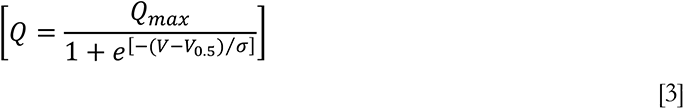

where *Q*_max_ is the maximal moveable charge, *V*_0.5_ is the voltage at which half of the charge is moved (that is, the midpoint of the sigmoidal), and *σ* = *kT*/*qδ* represents a slope factor, in which *q* is the elementary electronic charge, *k* is the Boltzmann constant, *T* is the absolute temperature, and *δ* is the fraction of electrical field over which the charge movement occurs.

Data were analyzed using Clampfit 10.7 (Molecular Devices). All figures and statistics were done with Origin 8.0 (OriginLab, Northampton, MA, USA).

### Solutions

The external control solution had the following composition: NaCl 98 mmol/L, MgCl2 1 mmol/L, CaCl2 1.8 mmol/L. For pH 5.5 and 6.5 the buffer solution Pipes 5 mmol/L was used; Hepes 5 mmol/L was used to obtain a pH 7.6. Final pH values were adjusted with HCl or NaOH. The substrate oligopeptide Gly-Gln (Sigma-Aldrich) was added at the indicated concentrations (from 3 µmol/L to 3 mmol/L) in the solutions with appropriate pH.

## Results

### Sequence and comparative analysis

Two Atlantic salmon *slc15a2* genes, namely *slc15a2a* (GenBank Acc. No. XM_014165384.1) and *slc15a2b* (GenBank Acc. No. XM_014173652.1), were identified in the GenBank database (**Table 1**). The two paralogues (*slc15a2a* and *slc15a2b*) encoded for proteins of 721 and 728 amino acids, respectively. PepT2a (GenBank Acc. No. XP_014020859.1) and PepT2b (GenBank Acc. No. XP_014029127.1) shared 93% similarity and 87% identity at the amino acid level. Comparative hydropathy analysis predicted 12 potential transmembrane domains with a large extracellular loop between transmembrane domains IX and X (**Fig. 1*A***). The structural key motif PTR2 family proton/oligopeptide symporter signature (orange diamonds; **Fig. 1*A***) was well conserved in Atlantic salmon PepT2 sequences, as well as the potential N-glycosylation site at the extracellular surface (brown circle; **Fig. 1*A***).

When the sequences of the Atlantic salmon PepT2a and PepT2b were analyzed for conserved functional residues (Killer *et al*., 2021), the proton and peptide binding sites (blue triangles; **Fig.1*A***), the amino acids that form the extracellular gate (green triangles; **Fig. 1*A***) and the intracellular salt bridge (red triangles; **Fig. 1*A***), as well as the amino acids involved in the subsequent steps of the transport cycle (1-5; **Fig. 1*A***), were conserved if compared not only to the mammalian PepT2 but also to mammalian and piscine PepT1 transporters. Notably, when compared with the rat (‘outward-facing open’ conformation) and human (‘inward-facing partially occluded’ conformation; substrate bound) PepT2 structures, Atlantic salmon PepT2a and PepT2b proteins both shared ∼55% identity with their mammalian counterparts (**Fig. 1*B***). Moreover, as emerged from the analysis of the superposed structures, (besides the large extracellular loop regions) little areas of ‘low consistency’ (red) amongst large areas of ‘high consistency’ (green) could be defined along the sequence, including some located within the transmembrane domains and other located at the basis of the extracellular loop stem (for details see **Fig. 1*B***).

Phylogenetic analysis of the putative Atlantic salmon PepT2a and PepT2b showed that these two proteins clustered into two distinct branches with their salmonid homologue sequences (**Fig. 2**). Northern pike PepT2 was the closest relative to the salmonid clade, and except for goldfish, which also underwent an additional WGD, only one PepT2-type protein was found in other teleost species.

**Figure 2.**
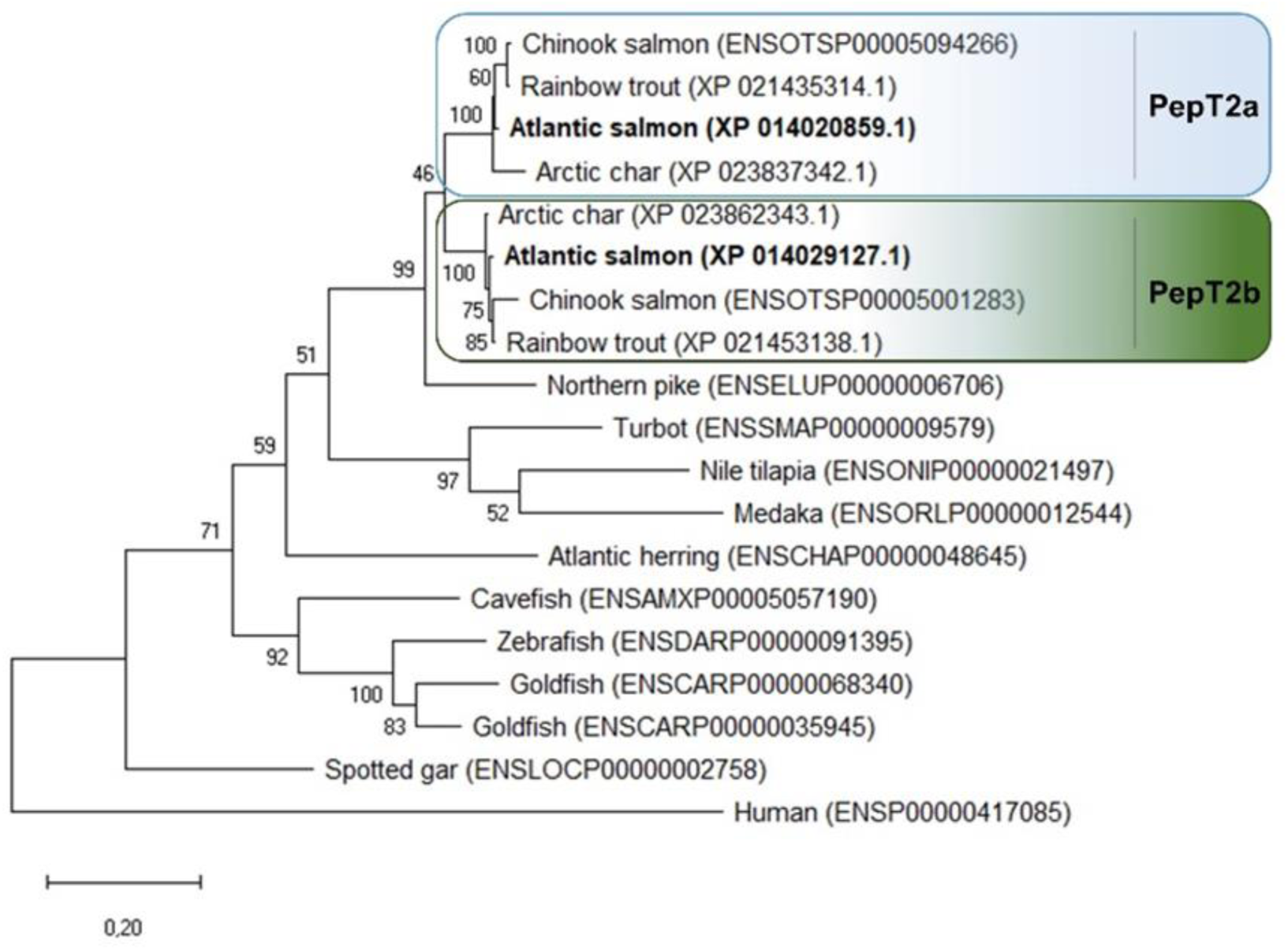
Phylogenetic relationship of fish and mammalian PepT2 based on predicted protein sequences. The (unrooted) phylogenetic tree was constructed based on deduced PepT2 amino acid sequences using the Maximum Likelihood (ML) method, 1000 bootstraps replicates, and JTT + G matrix-based model in MEGA X. The percentage of trees in which the associated taxa clustered together is shown next to the branches. Protein Acc. Nos. for GenBank or Ensembl databases are provided next to species common name.

The Atlantic salmon *slc15a2a* gene maps to chromosome ssa21 and the *slc15a2b* gene to chromosome ssa25. Both *slc15a2* paralogues share a conserved gene environment, with a total of 8 neighbouring genes upstream (out of 10) and 6 genes downstream (out of 10) being syntenic (**Fig. 3**). The conservation of synteny is particularly evident in the upstream region of *slc15a2*. Atlantic salmon *slc15a2a* upstream region shares 10 and 9 (out of 10) genes with the Northern pike and zebrafish homologue region, respectively, while for *slc15a2b* 8 genes are conserved in the Northern pike and zebrafish upstream homologue region. The genomic region downstream *slc15a2* is also conserved but to a lesser extent: salmon *slc15a2a* shares 5 and 3 genes with the Northern pike and zebrafish, respectively, while *slc15a2b* shares 8 genes with the Northern pike, but no genes are conserved in the downstream area with the zebrafish homologue region.

**Figure 3.**
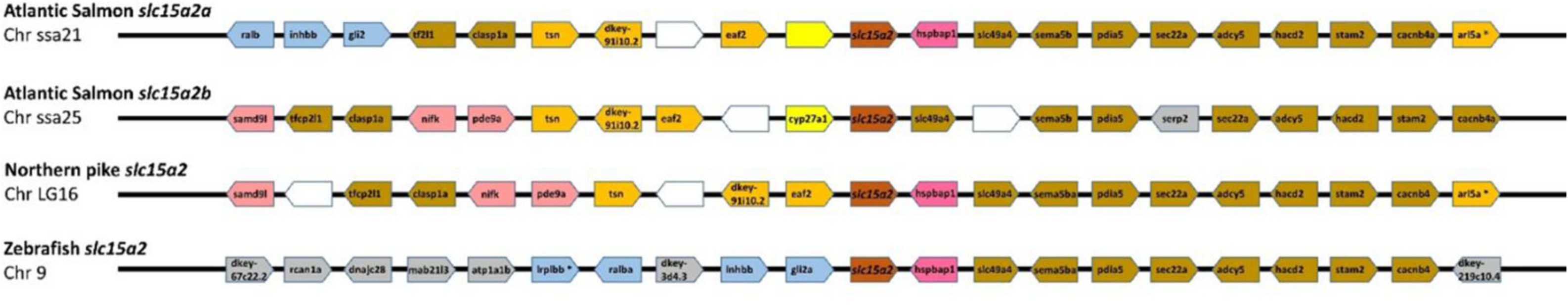
Synteny analysis of *slc15a2* genomic region in teleosts. From top *slc15a2a* and *slc15a2b* genes from Atlantic salmon (*Salmo salar*), *slc15a2* from Northern pike (*Esox lucius*) and zebrafish (*Danio rerio*). The chromosome (Chr) number is indicated below each species name. The central pentagons in dark orange indicate the *slc15a2* genes. For each *slc15a2* gene, 10 flanking genes upstream and downstream are represented by different coloured pentagons. Each colour identifies sets of orthologous genes based on the degree of conservation between species and between the chromosomes within species and white pentagons represent non-identified genes. The pentagons point in the direction of transcription and only protein-coding genes are indicated.

### Tissue distribution of Atlantic salmon slc15a2a and slc15a2b

Tissue expression analysis, focused on the Atlantic salmon head kidney and kidney (**Fig. 4*A***) and the alimentary canal and head tissues (**Fig. 4*B***), revealed that *slc15a2a* and *slc15a2b* has a different distribution profile. Atlantic salmon *slc15a2a* showed a wider tissue distribution profile with lower mRNA expression levels in the posterior stomach and higher levels in the brain and gills (**Fig. 4*B***). Interestingly, *slc15a2b* mRNA expression was mainly restricted to the gastrointestinal tract, specifically from pyloric caeca to the posterior hindgut, and excluding the stomach (**Fig. 4*B***). *slc15a2b* mRNA expression was abundant in the hindgut (**Fig. 4*B***), as well as in the kidney (**Fig. 4*A***).

**Figure 4.**
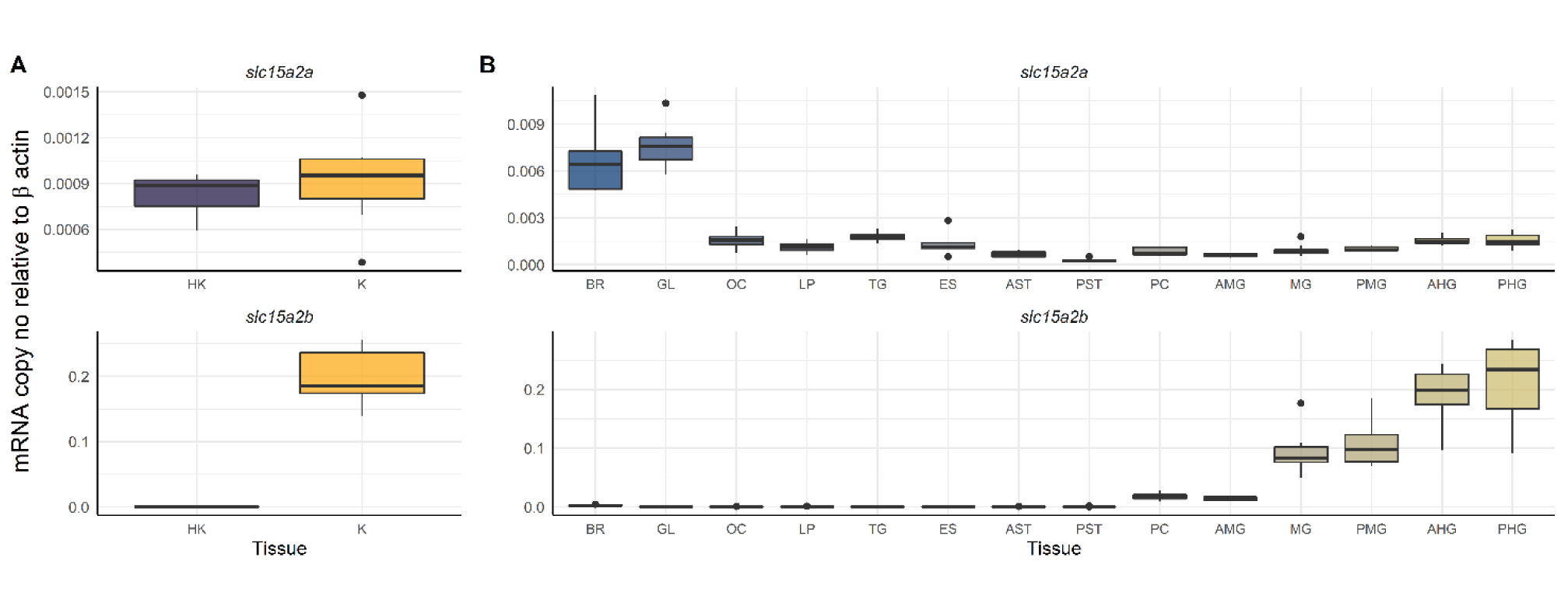
mRNA tissue distribution of *slc15a2a* and *slc15a2b* in head kidney and kidney in ***A*** and head tissues and along the gastrointestinal tract of the Atlantic salmon in ***B*** using quantitative RT-PCR. Results are shown as target *slc15a2* copy number per ng of total RNA normalized using *β-actin* copy number per ng of total RNA. The line in the boxplot indicates the median and boxes the 1^st^ to 3^rd^ quartile, whiskers mark variation outside 1^st^ and 3^rd^ quartile and dots the outliers (n = 8 for all tissues, except for HK, K, GL, OC, AMG, PMG and AHG where is n = 7). HK, head kidney; K, kidney; BR, brain; GL, gills; OC, olfactory cavity; TG, tongue; ES, esophagus; AST, anterior stomach; PST, posterior stomach; PC, pyloric caeca; AMG, anterior midgut; MG, midgut; PMG, posterior midgut; AHG, anterior hindgut; PHG, posterior hindgut.

### Transport currents

In **Fig. 5*A*** and ***C***, representative current traces recorded from oocytes expressing the two proteins are reported. At -60 mV both PepT2 paralogues are functional and elicit transport currents in the presence of 1 mmol/L Gly-Gln. The currents are larger at pH 5.5 and decrease with increasing pH, and even at pH 8.5 both proteins generate an inward current. PepT2a shows currents larger than PepT2b (**Fig. 5*A*- *D*** and **Fig. 6**). Moreover, for both PepT2a and PepT2b the current traces show an uncoupled current at acidic pH (5.5 and 6.5). These currents are visible in **Fig. 5*A*** and ***C*** by observing the position of the representative traces with respect of the dotted line, that is the zero current conventionally set at the holding potential at pH 7.6 for each transporter. This current is due to H^+^ entry at the acidic pH, mainly through the heterologously expressed transporters (leakage current), but also through endogenous oocyte channels.

**Figure 5.**
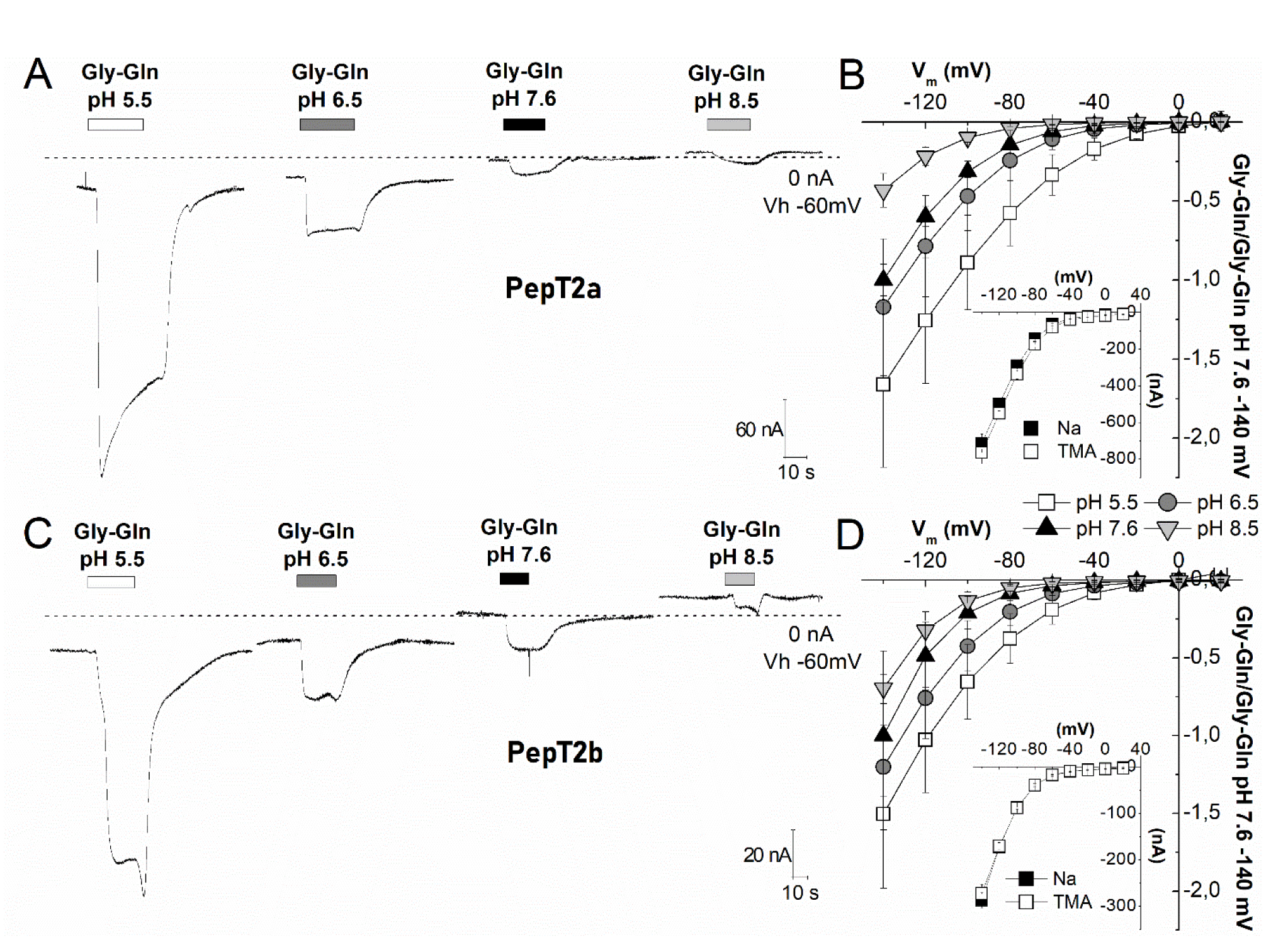
Transport of Gly-L-Gln in *Xenopus* oocytes expressing Atlantic salmon PepT2a (***A***, ***B***) and PepT2b (***C***, ***D***). In ***A***, ***C*** representative traces (1 mmol/L Gly-Gln in NaCl solution), the dashed line represents the baseline (conventionally fixed at the value in NaCl solution at pH 7.6). ***B***, ***D***, current/voltage relationships [the values are the mean (SD) of the current normalized to the mean value of the current at -140 mV and pH 7.6]; insets, currents at pH 6.5 in the presence of 98 mmol/L NaCl or tetraethylammonium chloride. Data are means (SD) from 4-24 oocytes from 1 to 5 batches.

**Figure 6.**
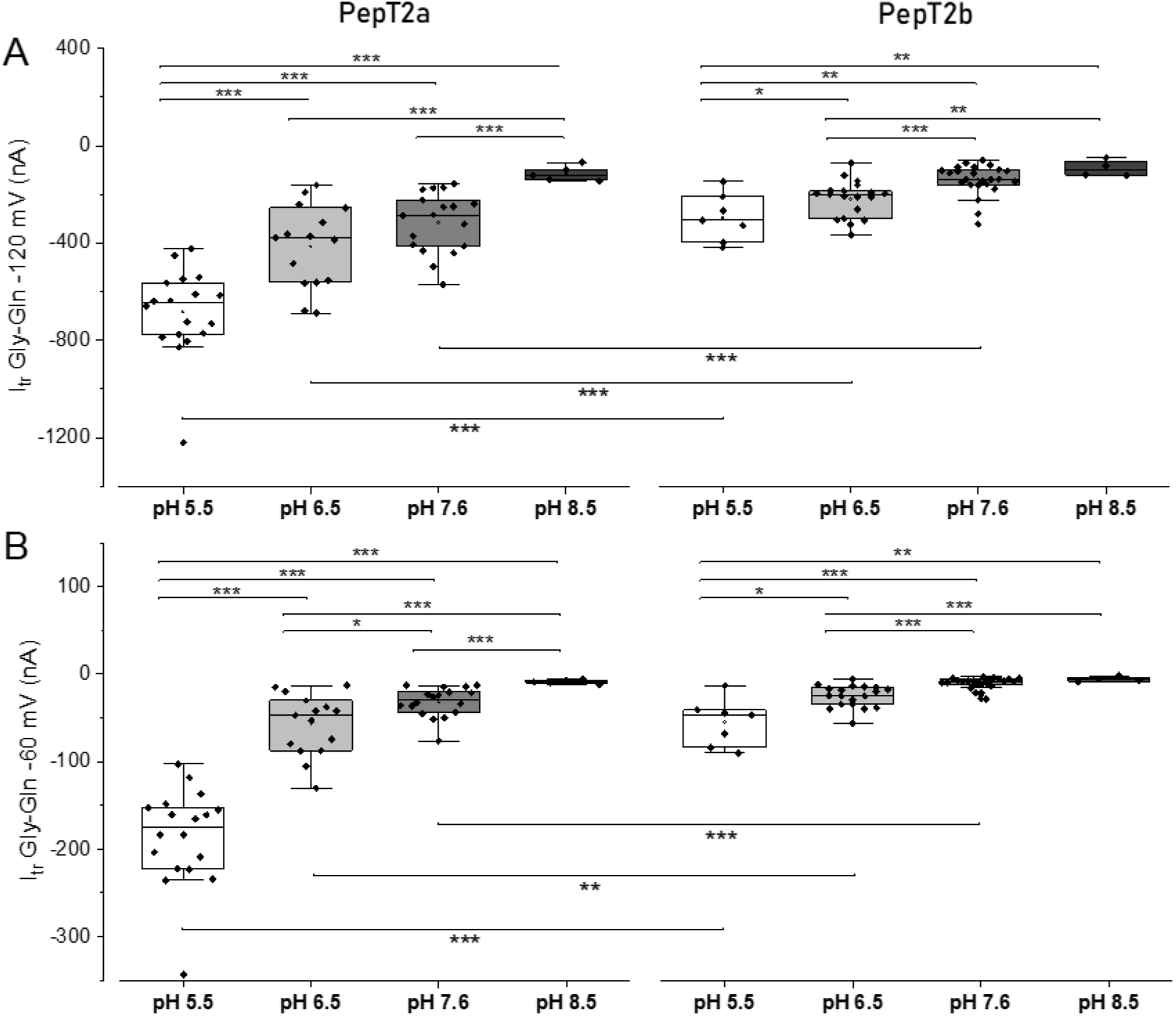
Box plots of the transport current values recorded as reported in Fig. 5, for PepT2a (left) and PepT2b (right) at -120 mV (***A***) and at -60 mV (***B***). Dots indicate the single oocyte transport current value. In the top part of each figure the statistical comparison between different pH conditions for each transporter is shown. In the bottom, the statistical comparison between transporters at the same pH (*Two sample t Test* or *Mann-Whitney U test*; * *p*<0.05, ** *p*<0.01 and *** *p*<0.001) is shown. Samples for box plots are the same as for Fig. 5. The detailed statistical values are reported in Statistical Summary Document.

The behaviour of PepT2a and PepT2b at different pH and different membrane voltages was investigated by measuring the transport associated current by applying the voltage pulse protocol described in Materials and Methods (see **Fig. 9** where the recorded traces are reported). The Atlantic salmon PepT2 proteins activity is pH- and voltage-dependent (**Fig 5*B*** and ***D***) and Na^+^-independent (**Fig. 5*B*** and ***D inset***). In the presence of the substrate, the decrease of external pH has a similar effect on the two transporters: i.e. the amplitude of the currents at pH 5.5 is the largest recorded in the voltage range from -140 mV to -20 mV (detailed by the box plot in **Fig. 6**). In PepT2a increasing the pH has more impact on the reduction of the transport associated current. In **Fig. 6*A*** and ***B***, the comparison of the transport currents (Itr) recorded at -120 mV and -60 mV at the indicated pH conditions confirmed that the amplitudes recorded are significantly different.

To understand the effect of pH on the transport it is essential to investigate the kinetic parameters in the different conditions. In **Fig. 7** the transport current-substrate concentration plot is reported for both PepT2 proteins at three pH values (5.5, 6.5 and 7.6). The relative maximal current (*I*_max_) and the Gly-L-Gln apparent affinity (1/*K*_0.5_) for each condition are given in **Table 3**.

**Table 3.**
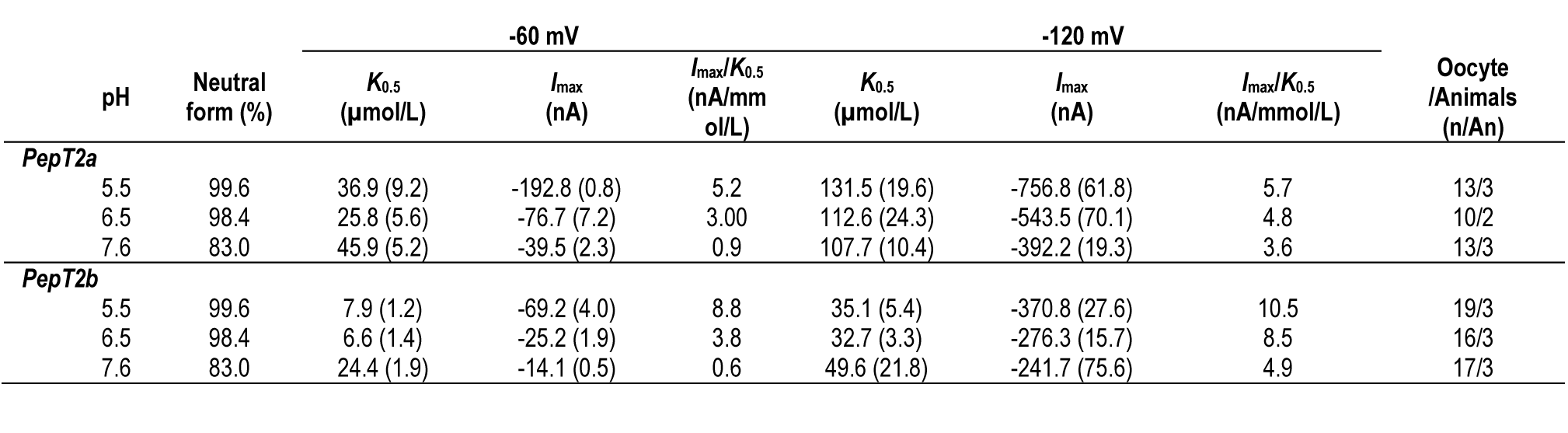
Kinetic parameters of the transport currents elicited by Gly-L-Gln in *Xenopus* oocytes. *K*_0.5_, *I*_max_ and *I*_max_/*K*_0.5_ values from **Fig. 8** were reported in the table for the -60 mV and -120 mV. Kinetic parameters were calculated by least-square fit to the Michaelis-Menten equation [1] and the values are expressed as mean (±SE). *I*_max_/*K*_0.5_, transport efficiency.

**Figure 7.**
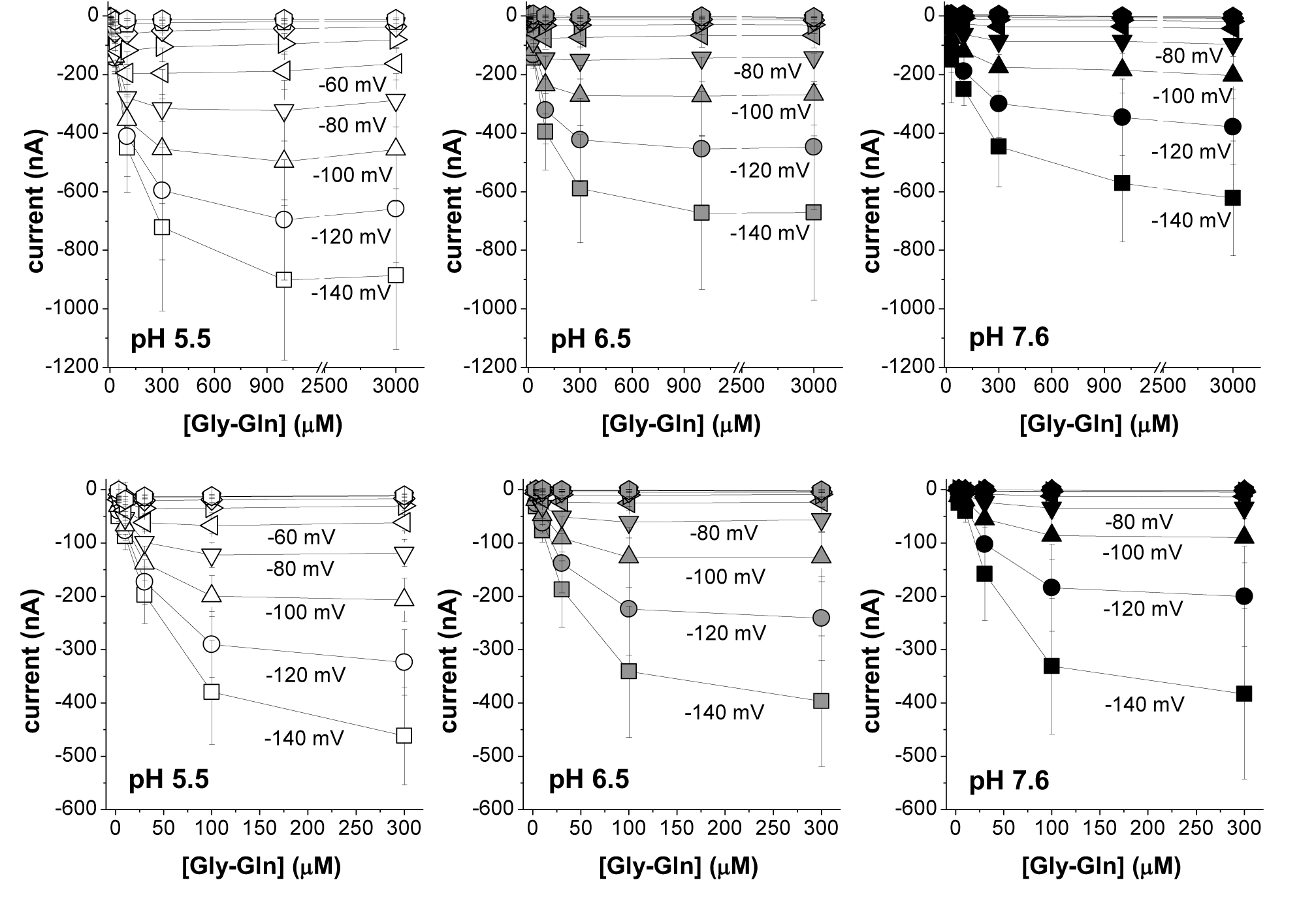
Current *vs.* substrate concentration (*I*/*S*) relationships at the indicated voltages for Atlantic salmon PepT2a in ***A***, ***B*** and ***C*** and for PepT2b in ***D***, ***E*** and ***F***. The mean value of the currents at each concentration (from 3 µmol/L to 3 mmol/L for PepT2a and from 3 µmol/L to 300 µmol/L for PepT2b) are plotted at the indicated voltage and pH. Data are means (SD) from 10-19 oocytes, obtained from 2 to 3 batches.

The *I*_max_, *K*_0.5_ and transport efficiency are reported for PepT2a in **Fig. 8*A***, ***B*** and ***C*** and for PepT2b in ***D***, ***E*** and ***F***. These parameters allow to evaluate the functional characteristics and the differences between the two proteins. As suggested by the *I*_max_ *vs.* voltage (*I*_max_/*V*) graphs in **Fig. 8*A*** and ***D***, the *I*_max_ of the two PepT2 transporters is voltage dependent, and the current amplitudes increase from 0 to -140 mV for both proteins and at all tested pH. Rising the extracellular pH from 5.5 to 7.6 results in a pronounced decrease in the maximal transport currents of ∼5-fold at -60 mV [i.e. from -192.7 (20.8) nA at pH 5.5 to -39.4 (2.3) nA at pH 7.6 for PepT2a and from -69.2 (4.0) nA at pH 5.5 to -14.1 (0.5) nA at pH 7.6 for PepT2b]. Notably, the *I*_max_ recorded for PepT2a at -60 mV is ∼3-fold higher than the *I*_max_ current recorded at the same membrane voltage for PepT2b, for all tested pH.

**Figure 8.**
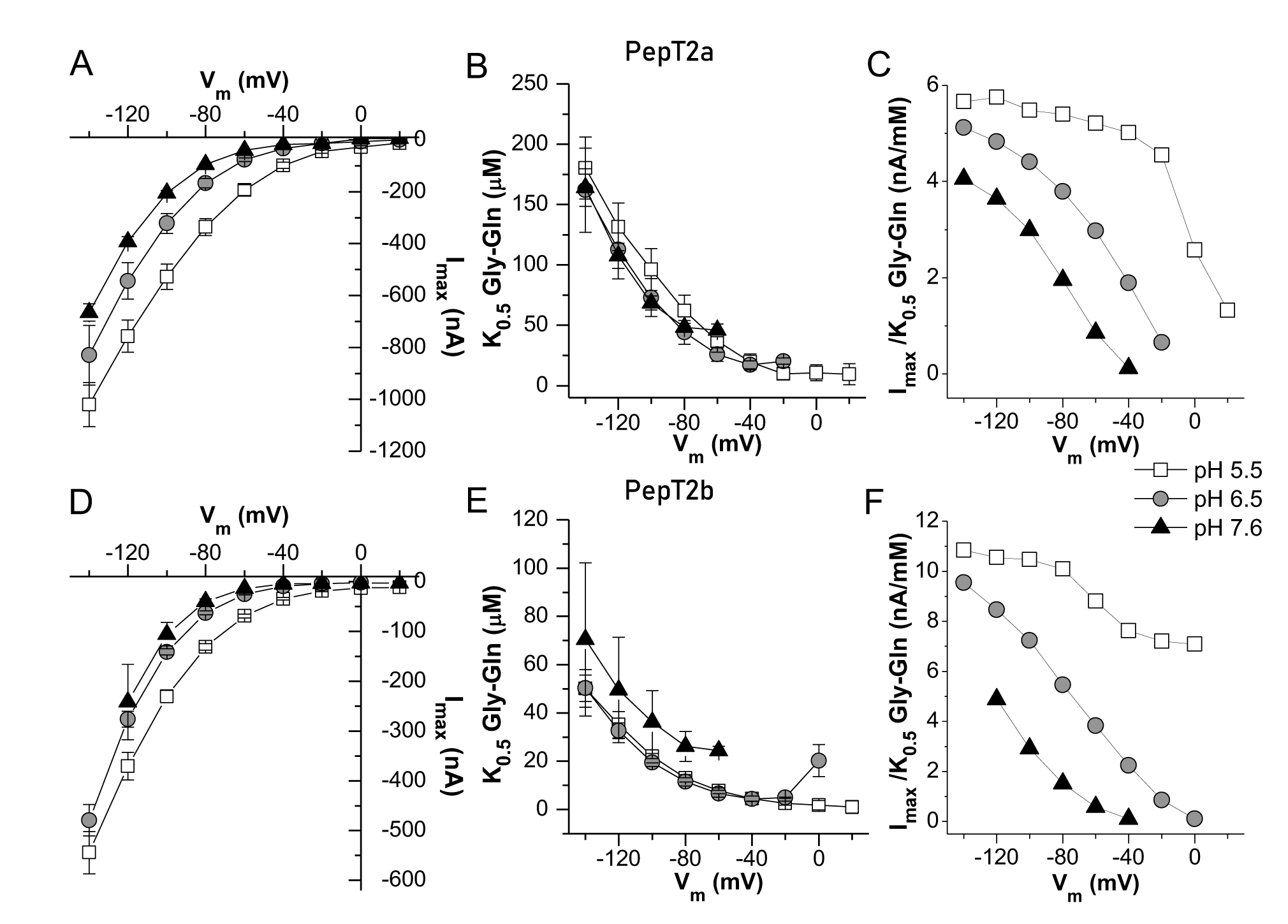
Dose response analysis: *I*_max_, *K*_0.5_ and transport efficiency of Atlantic salmon PepT2a in ***A***, ***B*** and ***C*** and PepT2b in ***D***, ***E*** and ***F***. The current values evaluated in the presence of increasing concentrations of Gly-L-Gln under each tested voltage (Fig. 3) were subsequently fitted with the equation [1] to obtain the relative maximal current (*I*_max_) in ***A*** and ***D***, the *K*_0.5_ in ***B*** and ***E***, i.e. the substrate concentration that elicits half of the maximal current (*I*_max_), and the transport efficiency in ***C*** and ***F***, evaluated as the ratio *I*_max_/*K*_0.5_ under each membrane potential and pH condition.

**Figure 8*B*** and ***E*** report the *K*_0.5_/*V* curves for PepT2a and PepT2b, respectively. For both transporters at all external pH conditions, *K*_0.5_ values exhibit a peculiar dependence on membrane potentials with an increase of *K*_0.5_ values from -60 mV to -140 mV. *K*_0.5_ values are only slightly modified by the acidification of the bathing solution: at -60 mV, *K*_0.5_ is in the range of tens of μmoles per liter for all the pH for both transporters. However, PepT2b has higher affinity in all the conditions tested for Gly-L-Gln if compared to PepT2a.

As most of solute carriers, the larger impact on the substrate apparent affinity is due to the hyperpolarization of the membrane voltage that increases the driving force and supports the entrance of substrate and H^+^. The efficiency of the transport reflects the two parameters, and it is maximal, as expected, at pH 5.5 at most hyperpolarizing conditions (**Fig. 8*C*** and ***F***). Due to the higher affinity of PepT2b, the efficiency is slightly higher in this transporter and the best working condition is at -120 mV at pH 5.5.

For the experiment at pH 7.6, for both PepT2-type proteins the currents recorded at membrane voltages higher than -60 mV were too low, and the *K*_0.5_ or *I*_max_ obtained were not reliable and, thus, not considered in the analysis.

### Presteady-state currents

When rectangular voltage jumps (pulses from -140 to +20 mV in 20 mV increments) protocol is applied to oocytes expressing a transporter, the elicited currents in the absence and presence of the substrate can be a tool for biophysical investigations on the steps of the transport cycle.

Slow transient currents, related to the presence of the transporter in the plasma membrane, were only observed in oocytes expressing the two PepT2-type proteins, and almost completely abolished by addition of saturating Gly-L-Gln (1 and 0.3 mmol/L for PepT2a and PepT2b, respectively) (**Fig. 9**).

**Figure 9.**
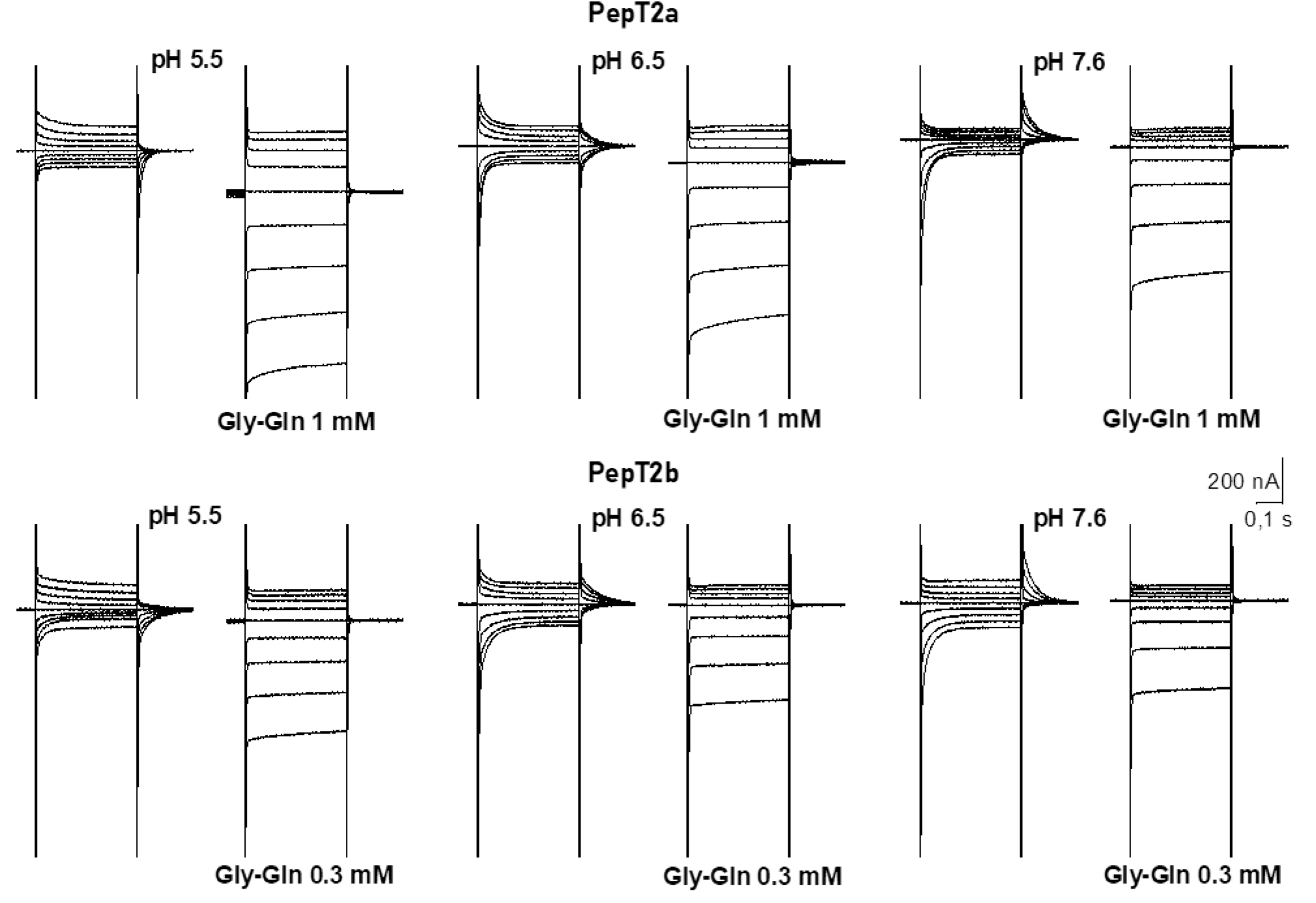
Representative traces of currents elicited by voltage steps protocol applied to *Xenopus* oocytes expressing the Atlantic salmon PepT2 transporters, PepT2a (top) and PepT2b (bottom). The voltage pulses were in the range -140 to +20 mV. At all tested pH, the presence of saturating concentrations of Gly-L-Gln (1 mmol/L for PepT2a and 0.3 mmol/L for PepT2b) eliminates the transporter transient component (*I*_PSS_) and produces large inwardly directed steady-state currents that raise increasing the proton chemical gradient and in hyperpolarization.

The behaviour of the currents reported in **Fig. 9** shows that the pH of the bathing solution affected the current kinetics in both transporters. At pH 5.5, in the absence of substrate, slow transient currents are mostly present in response to depolarizing pulses while at pH 7.6 they are principally present in response to hyperpolarizing pulses; at pH 6.5, the transient currents are symmetrically arranged around the holding potential and are similar for amplitude and decay time (**Fig. 9** and **Fig. 10*A*** and ***C***) for hyperpolarizing and depolarizing voltages. This behaviour is more pronounced in PepT2a, where the symmetry at pH 6.5 is almost perfect around the holding voltages.

**Figure 10.**
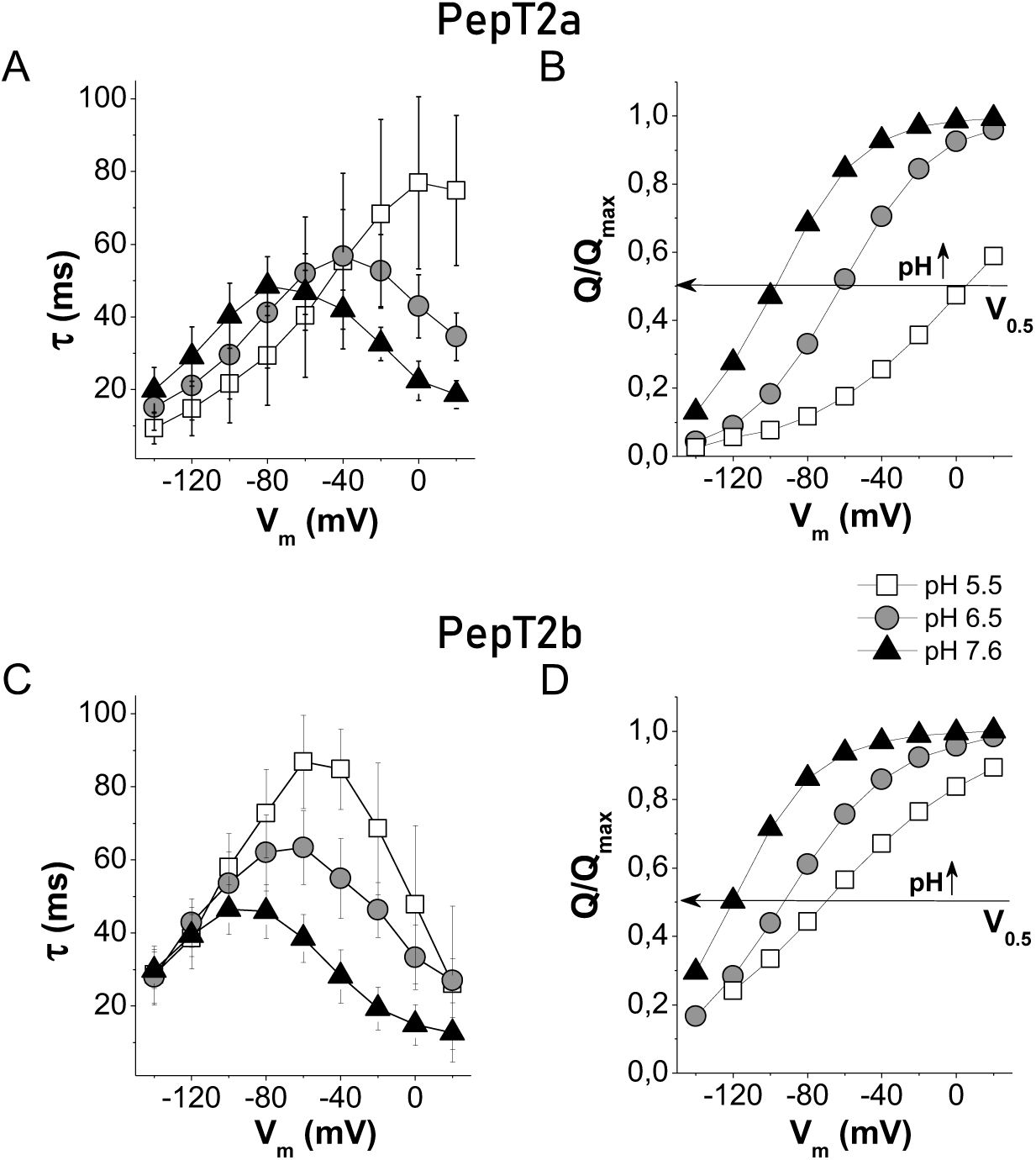
The time constant of decay (τ) and normalized displaced charge (*Q*/*Q*_max_) for Atlantic salmon PepT2 transporters, PepT2a in ***A*** and ***B*** and PepT2b in ***C*** and ***D.*** *τ*/*V* in ***A*** and ***C*** and (*Q*/*Q*_max_)/*V* in ***B*** and ***D***. The sample sizes for *τ*/*V* and *Q*/*V* for each condition were between 14 and 22 oocytes, from 2 to 3 batches. The *τ*/*V* values are the mean (SD). The *Q*/*V* values are the mean normalized against the maximal moveable charge *Q*_max_ values obtained by fitting using the equation [3]. The unitary charge level was set to the saturation value at positive potential. In ***B*** and ***D***, the horizontal arrow intersects each curve at half completion of charge movement corresponding to *V*_0.5_ given by the Boltzmann equation [3].

To study the *I*_PSS_ component elicited by the voltage jumps, from the traces reported in **Fig. 9** the *I*_PSS_ were separated from the capacitive current of the oocytes as described in Material and Methods (see double-exponential function, equation [2]), assuming that the slow component represents the *I*_PSS_ (Sangaletti *et al*., 2009). The currents were then analyzed to obtain the biophysical parameters of the transport.

The decay time constant is measured at each potential and plotted (*τ*/*V*). The area underneath the *I*_PSS_ currents correspond to the amounts of charges (*Q*), intrinsic and extrinsic, moved in the membrane electric field (Peres *et al*., 2004a; Bossi *et al*., 2011; Renna *et al*., 2011b).

The decay time (*τ)* and amounts of charges (*Q*) as a function of potential (*V*), *τ*/*V* and *Q*/*V*, respectively curves, at each tested pH are reported in **Fig. 10**. Like other peptide transporters, such as mammalian and fish PepT1 (Renna *et al*., 2011a; Vacca *et al*., 2019) and rat PepT2 (Chen *et al*., 1999), the *τ*/*V* curves of both PepT2 appeared bell-shaped (**Fig. 10*A*** and ***C***). Data on *τ*_max_ at different pH for the two transporters are reported in **Table 4**.

**Table 4.** The maximal value of the decay time constant (*τ*_max_) and Boltzmann equation parameters of Atlantic salmon PepT2 proteins (PepT2a and PepT2b) calculated at pH 5.5, 6.5 and 7.6. *Q*hyp and *Q*dep represent the amount of displaced charge at hyperpolarizing and depolarizing limits, *Q*_max_ (= *Q*dep- *Q*hyp ) is the maximal moveable charge, *V*_0.5_ is the voltage at which half of the charge is moved, *σ* represents a slope factor of the sigmoidal curve (Renna *et al*., 2011b). Parameters were calculated by nonlinear fit to the Boltzmann equation [3] using Origin 8.0 from the curves of Fig. 10. The number of oocytes for each condition were between 14 and 22, from 2 to 3 batches.

| pH | PepT2a |  |  | PepT2b |  |  |
| --- | --- | --- | --- | --- | --- | --- |
|  | 5.5 | 6.5 | 7.6 | 5.5 | 6.5 | 7.6 |
| $\tau_{\text{max}}$ (ms) | 76.8 ± 6.3 | 56.7 ± 3.6 | 48.5 ± 1.7 | 86.9 ± 3.1 | 63.4 ± 2.3 | 46.5 ± 3.1 |
| $Q_{\text{hyp}}$ (nC) | -2.82 ± 0.12 | -7.35 ± 0.05 | -11.8 ± 0.1 | -9.19 ± 0.37 | -14.0 ± 0.2 | -17.5 ± 0.4 |
| $Q_{\text{dep}}$ (nC) | 13.2 ± 1.2 | 6.80 ± 0.05 | 2.20 ± 0.03 | 7.07 ± 0.18 | 4.49 ± 0.06 | 1.21 ± 0.03 |
| $Q_{\text{max}}$ (nC) | 16.1 ± 1.3 | 14.2 ± 0.1 | 14.0 ± 0.2 | 16.3 ± 0.6 | 18.5 ± 0.3 | 18.8 ± 0.5 |
| $V_{0.5}$ (mV) | 4.85 ± 5.90 | -62.1 ± 0.3 | -97.6 ± 0.5 | -70.7 ± 1.5 | -93.0 ± 0.7 | -120.5 ± 1.2 |
| $\sigma$ (mV) | 42.1 ± 2.6 | 25.1 ± 0.3 | 22.5 ± 0.4 | 42.9 ± 1.7 | 29.2 ± 0.6 | 22.4 ± 0.5 |

To better appreciate the pH effect on *Q*/*V* curves, the mean values of *Q*_on_-off fitted with the Boltzmann equation [3] and normalized against the maximal moved charge (*Q*_max_) setting the unitary charge level to the saturation value at positive potential (Forlani *et al*., 2001) are reported in **Fig. 10*B*** and **Fig. 10*D***.

In both transporters, the sigmoidal shape of the charge *vs.* voltage curves showed a clear pH dependence with an inflexion point (**Fig. 10*B*** and ***D***), corresponding to the half of the charge translocation (*Qmax*/2), shifted to more negative values of membrane potential (*V*_0.5_ values in **Table 4**) as pH increased from 5.5 to 7.6. At pH 7.6 and in depolarization condition (from -20 mV to +20 mV) the *Q*/*V* curves showed that PepT2 transporters reached the saturation of the amount of displaced charge. Conversely, charge saturation at negative potential is reached only by PepT2a, at pH 5.5, and in this case, at +20 mV the amount of moved-charge is only close to half *Q*_max_ quite far from the predicted saturation value at depolarizing value (**Fig. 10*B***). PepT2b at pH 5.5 presents only the central part of the sigmoid curves, a condition that agrees with the shape of the *τ*/*V* curves (**Fig. 10*D***).

According to the two-state system representation of the charge movement process (see cartoon in **Fig. 11**), the parameters obtained by fitting of charge *vs*. voltage curves (*Q*/*V*) with the Boltzmann function (*Q*_in_ and *Q*_max_) and the time decay constant *vs*. voltage curves (*τ*/*V*) can be used to determine the unidirectional inward, and outward rate constants of the charge movement (Renna *et al*., 2011b).

**Figure 11.**
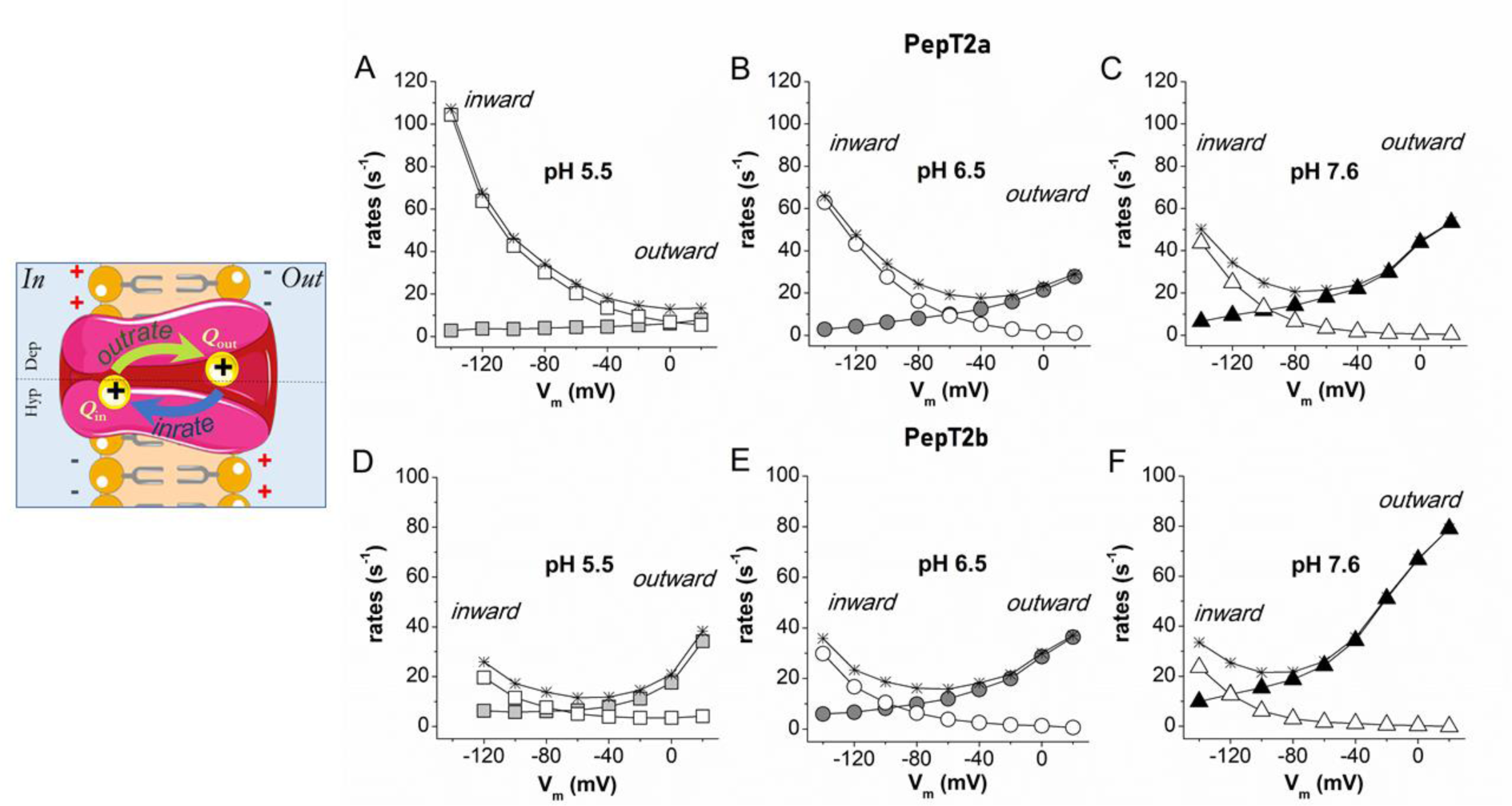
The unidirectional rate constants. Graphical representation of a two-state system (left). Inward (open symbols) and outward (solid symbols) rate constants of the intramembrane charge movement of Atlantic salmon PepT2 transporters, PepT2a in ***A***, ***B*** and ***C*** and PepT2b in ***D***, ***E*** and ***F***. The unidirectional rates, plotted as function of membrane potentials, were calculated from *τ*/*V* (and (*Q*/*Q*_max_)/*V* (in Fig. 10) at the different pH. The rate (1/τ) of the two transporters are plotted as stars.

The charge movement process can be described with a simple reaction where the outrate and the inrate are the unidirectional rate constants:

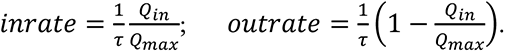

*Q*_in_ is the amount of charge located near the inner side of the membrane electrical field and *Q*out is the amount of charge displaced toward the outer membrane margin.

In **Fig. 11**, the voltage and pH dependence of the unidirectional rate constants for Atlantic salmon PepT2a and PepT2b transporters are reported. Assuming a positive mobile charge, the outward rates increase when the inner side of the cellular membrane presents a positive voltage and decreases to zero for membrane voltage values lower than -100/-140 mV. Conversely, the inward rates increase and decrease, respectively, with hyperpolarization and depolarization of the membrane potential.

Within the voltage range tested, both the inward and outward rates of PepT2 proteins showed a change in the voltage dependence with the changes of pH (from 5.5 to 7.6). The inward rates are more affected by voltage than the outward rates in PepT2a than PepT2b (Fig. 11). Little or no change appears to occur for the inward rate for PepT2b (Fig. 11D, E), instead a great chaghes is visible in PepT2a at pH 5.5. The outward rate instead was more influenced by the voltage at pH 7.6 in Pept2b.

## Discussion

In this study, two *slc15a2* genes, named *slc15a2a* and *slc15a2b*, were examined in the Atlantic salmon. Comparison of sequence, structure and gene environment strongly suggests that these two high-affinity/low-capacity transporters have evolved from the WGD event that salmonids experienced 94 million years ago (Macqueen & Johnston, 2014; Lien *et al*., 2016), where many of the duplicated genes were retained as functional copies. This finding is corroborated by the parallel observation that all other teleost species analyzed in this study have only one *slc15a2* gene, with the exception of the salmonids and the goldfish (a cyprinid), which genome has also undergone an additional WGD event (Kuang *et al*., 2016) and also exhibits two *slc15a2* genes. Noteworthy, in a very recent Slc15 family-centered study in the common carp (*Cyprinus carpio*) genome, another piece of information has emerged on the presence of two genes encoding for PepT2-type transporters in one genome, with the *slc15a2a-1* and *slc15a2-2* genes well-described in this cyprinid (Dong *et al*., 2020). In the Atlantic salmon, genes originating from the salmonid specific WGD have often been shown to take on new functions (neo-functionalization) or sub-functions (sub-functionalization) of their duplicates (Lien *et al*., 2016). It is very likely that sub-functionalization has occurred for the Atlantic salmon *slc15a2a* and *slc15a2b*, too. In fact: i) the Atlantic salmon PepT2a and PepT2b transporters, generated by *slc15a2a* and *slc15a2b* respectively, are conserved, sharing 87% identity at the amino acid level, and exhibiting all the essential PepT2-type functional motifs; ii) based on the gene expression data, after the gene duplication event functional divergence may have occurred, and the two identified genes, *slc15a2a* and *slc15a2b*, may have acquired some tissue-specific functions. In this respect, the expression of *slc15a2b*, particularly the very high levels observed in the Atlantic salmon kidney (see **Fig. 4*A***), is consistent with the well-known primary function of PepT2 in reabsorbing di/tripeptides from the ultrafiltrate [see e.g. (Boll *et al*., 1996; Rubio-Aliaga *et al*., 2000; Romano *et al*., 2006)]. Moreover, similarly to what previously reported in a variety of teleost fish species (summarized in **Table 1**), the mRNA expression of *slc15a2b* in the gastrointestinal tract of the Atlantic salmon is mainly confined to the mid-to-distal intestine, a pattern that has been previously observed in the same species (Del Vecchio *et al*., 2021) and also found in e.g. the rabbit alimentary canal (Döring *et al*., 1998). The presence of *slc15a2b* in the distal area of the intestine is most probably related to the enterocytes that secure absorption of small peptides, although the fact that a PepT2-type transporter is also expressed by neurons of the enteric nervous system (Rühl *et al*., 2005) cannot be ruled out a priori. In contrast, in the Atlantic salmon, the *slc15a2a* paralogue exhibits a widespread tissue distribution, in terms of the presence of mRNA, but it is most abundant in the brain and, notably, in the gills. The expression of a PepT2 in the brain, operating as a di/tripeptide uptake system, has been observed in mammals (Wada *et al*., 2005; Biegel *et al*., 2006; Kamal *et al*., 2008; Keep & Smith, 2011; Smith *et al*., 2013; Viennois *et al*., 2018) as well as in teleost fish species, e.g., zebrafish (Romano *et al*., 2006) and Mozambique tilapia (Con *et al*., 2019). But, noteworthy, the identification of PepT2 mRNA transcripts in the gills opens to the hypothesis that a di/tripeptide transporter might also be operating in such an epithelial tissue moving di/tripeptides directly from the aquatic environment to the blood and/or vice versa.

The fact that key amino acids involved in substrate recognition and transport are well conserved between Atlantic salmon and mammalian (rat and human) PepT2 proteins (**Fig. 1*A***), and that both salmon PepT2a and PepT2b protein structures share about 55% identity with the mammalian homologue (**Fig. 1*B***), indicates that their function is conserved across species. Indeed, both PepT2a and PepT2b in the presence of dipeptides (Gly-L-Gln) elicit inward currents, as it happens for the other Slc15 members (Saito *et al*., 1996; Chen *et al*., 1999; Rubio-Aliaga *et al*., 2000; Terada *et al*., 2000; Biegel *et al*., 2006; Romano *et al*., 2006; Verri *et al*., 2017). These currents are pH-dependent and Na^+^-independent, with inward coupled currents being recorded even when Gly-L-Gln is perfused at pH 8.5. However, the transport current is three times higher in PepT2a compared to PepT2b and at pH 5.5 the current rapidly decreases after the first seconds of substrate perfusion (**Fig. 5**). This phenomenon is not new in solute carriers (Mackenzie *et al*., 1996; Eskandari *et al*., 1997; Mackenzie *et al*., 1998), and in PepT2 transporters (Chen *et al*., 1999). It is observed with cotransporters expressed in oocytes, mainly at substrate concentrations above the apparent *K*_0.5_. The decreases in current after the peak can be due to decreased H^+^ concentrations at the immediate proximity of the extracellular membrane or to the increase of intracellular substrate or H^+^ accumulation, in analogy to explanations for PepT1 (Kottra & Daniel, 2001; Kottra *et al*., 2002; Kottra *et al*., 2009; Bossi *et al*., 2011) and PepT2 (Chen *et al*., 1999). For PepT2b, however, a different behaviour is observed. In this transporter, the removal of 1 mmol/L Gly-Gln at the end of the perfusion time induces a rapid and fast increasing current at pH 5.5, suggesting that substrate concentration at 1 mmol/L, which is a concentration higher than the saturating value of 300 μM, reduced the final transport current. The behaviour is related to the accumulation of substrate inside the cell and to the high affinity of the transporter for the substrate. In this case, the net flux at very high substrate concentration results lower than the flux at the concentration proximal to saturation value (Mertl *et al*., 2008; Renna *et al*., 2011a; Bosdriesz *et al*., 2018).

Dose-current experiments at different pH and voltage confirm that the two Atlantic salmon PepT2 proteins behave as high-affinity/low-capacity transporters. The ‘canonical’ renal transporter PepT2b shows a higher affinity and smaller current if compared to the ‘brain/gills’ PepT2a. When the apparent affinity (1/*K*_0.5_) and *I*_max_ are determined and plotted *vs.* voltage at different pH, other functional characteristics distinguish the two Atlantic salmon proteins. Atlantic salmon PepT2a and PepT2b tested in the same conditions, using the same protocol and substrate, have specific transport characteristics which are underlined by the detailed biophysical and kinetic analysis presented in **Fig. 8**. The data of PepT2b transporter are very similar to the data collected for the zebrafish (Romano *et al*., 2006) and recall the rabbit PepT2 (Amasheh *et al*., 1997), but with less marked pH differences. In fact, in Atlantic salmon PepT2b, the reduction of H^+^ is balanced by the hyperpolarizing voltages [see comment in: (Sala-Rabanal *et al*., 2008)].

Many solute carriers have charged residues located in the transmembrane helices. When the membrane voltage is rapidly changed across the membrane (voltage jumps) some charges of these residues turn to new equilibrium positions, giving rise to *I*_PSS_ currents (Lester *et al*., 1996; Mager *et al*., 1996; Loo *et al*., 1998; Peres *et al*., 2004b). These currents (*I*_PSS_) disappear when the new equilibrium is achieved. Furthermore, binding or unbinding of substrate or more often of the driving ions is voltage-dependent and can contribute to the generation of the *I*_PSS_ currents (Bossi *et al*., 2002; Fesce *et al*., 2002; Bossi *et al*., 2012; Cherubino *et al*., 2012). In this respect, the presteady-state currents (*I*_PSS_) analyses give a unique opportunity to collect information on the kinetic properties and the transport cycle of the Atlantic salmon PepT2 transporters. In contrast to PepT1 transporters, which have deeply been characterized in both mammals and teleosts (Mackenzie *et al*., 1996; Nussberger *et al*., 1997; Sangaletti *et al*., 2009; Renna *et al*., 2011b), data on presteady-state currents (*I*_PSS_) of PepT2 are lacking. Only two papers (Chen *et al*., 1999; Sala-Rabanal *et al*., 2008) analyzed few conditions, limited to rat and human transporters, and information about the relation of charge movements and transport processes in PepT2 is otherwise completely missing from other species and fish. As previously reported for oocytes expressing the rat (Chen *et al*., 1999) and human (Sala-Rabanal *et al*., 2008) PepT2, Atlantic salmon PepT2a- and PepT2b-injected oocytes do show transient currents after step changes in membrane potential. Unlike most transporters in which *I*_PSS_ currents have been analyzed, in mammalian PepT2 two peculiar effects were reported. First, both Chen (Chen *et al*., 1999) and Sala-Rabanal (Sala-Rabanal *et al*., 2008) report that the presence of *I*_PSS_ is increased at acidic pH in the presence of non-saturating substrate concentrations (lower than *K*_0.5_) for human transporter and of saturating substrate concentrations for the rat transporter. Moreover, for the human transporter a discrepancy of the *Q*_on_ and *Q*_off_ has also been observed, according to the duration of the voltage step. Instead, the behaviour of both salmon PepT2 *I*_PSS_ and the relative parameter is more similar to the “classical” model and to that reported for PepT1. The *I*_PSS_ were larger in the absence of substrate at all pH tested and *Q*_on_ and *Q*_off_ were always similar. The effect of varying the external H^+^ concentration on PepT2 *I*_PSS_ kinetics is emphasized by the shift toward more negative potentials of both the maximal time constant (*τ*_max_) and the half of the moved maximal charge (*Q*_max_*/*2), in response to increasing pH of the external medium. Accordingly, the same effect of the pH is observed on the shift of the crossing point between the inward and outward rates. In Atlantic salmon PepT2a, the saturation of the amount of displaced charge at more hyperpolarizing potentials at pH 5.5 is coherent with the high acceleration of the inward rate. Conversely, for both Atlantic salmon PepT2, at pH 7.6 the saturation of the moved charge at depolarizing potentials is reached and it associates to an acceleration of the outward rate. This effect is more evident in Atlantic salmon PepT2b.

All together, these data confirm that for the two Atlantic salmon PepT2 transporters the polarity of *I*_PSS_ kinetics and the magnitude of the charge movements is strictly dependent on the H^+^ electrochemical potential across the plasma membrane (Mager *et al*., 1998).

The comparison between the *V*_0.5_ values of PepT2a and PepT2b reveals marked differences at the same pH with *V*_0.5_ of Atlantic salmon PepT2a right-shifted (**Table 4**). The slope factor (*σ*) is instead similar at each pH showing that the two PepT2 transporters exhibit a similar fraction of the electrical field where the charge movement occurs. It is also important to highlight that the *Q*_max_ showed similar values in all the conditions (**Table 4**). Considering that the *I*_PSS_ are mainly due to intrinsic charges of the protein that are movable, the fact that *Q*_max_ does not change in the tested conditions suggests that the expression on the membrane of the *Xenopus* oocytes of the two transporter is similar (Mager *et al*., 1993; Forster *et al*., 2006).

To summarize, our experimental data suggest that for both Atlantic salmon PepT2 transporters i) the maximal charge displacement (*Q*_max_) is independent of external proton concentration (**Table 4**); ii) the charge movement in the on-response (*Q*_on_) is balanced by the charge moved in the off transition (*Q*_off_) at all potential jumps from the holding voltage; iii) the addition of substrate at saturating concentration eliminates the *I*_PSS_ for all tested pH conditions and produces large inwardly directed steady-state transport currents; and iv) acidification causes a positive shift in the voltage dependence of both presteady-state and steady-state related parameters. Based on all four observations listed above qualitatively kinetic models can be hypothesized, in which in the empty transporters negatively charged and protonated “trapping” states are introduced to account for the voltage shift produced by pH. Indeed, the kinetic schemes proposed by (Sala-Rabanal *et al*., 2008) and (Chen *et al*., 1999) can qualitatively account for the effect of the pH on the unidirectional rate constant shown in **Fig. 11** and consequently of the *τ*/*V* and *Q*/*V* curves of **Fig. 10**. These schemes are shown in **Fig. 12** and include a voltage-independent and fast binding of a second proton (transition 2 →7 in **Fig. 12*A*** or II→III in **Fig. 12*B*)**. As detailed in (Mager *et al*., 1998) for the neurotransmitter transporter GAT1, sequential binding of a second driving ion would reduce the outward rate constant of charge movement, an effect observed in GAT1 with Na^+^ ions and in both PepT2a and PepT2b with H^+^. This effect is much more significant in rat and human PepT2 transporters. Conversely, the transition 1←2 or I←II represented in **Fig. 12** dependent on external protons will speed up the inward rate, particularly for PepT2a.

**Figure 12.**
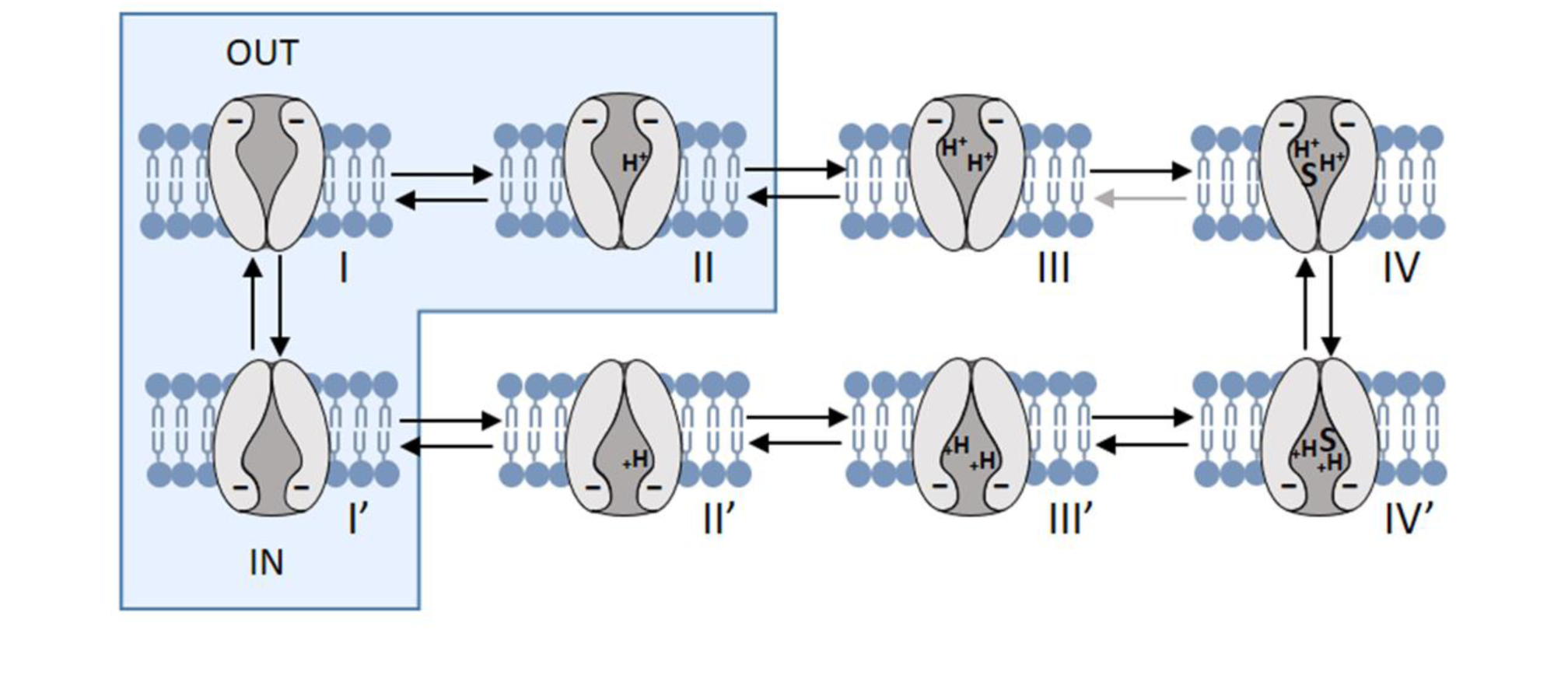
Representation of two kinetic models for PepT2 transporter. Each configuration represents a state, from I to IV (outside) and from I’ to IV’ (inside). The presteady-state currents arise from the redistribution of the negatively charged transporter between states I’, I and II (blue boxes). The binding of the first external proton to state I accelerates the inward rate constant, while the binding of a second proton to state II leads to state III, effectively slowing down the outward rate. Transitions between I’, I and II are voltage-dependent. [S] represents the binding of the substrate. Modified from (Chen *et al*., 1999; Sala-Rabanal *et al*., 2008).

The different behaviour at acidic pH between piscine and mammalian transporter could be related to the potency of the “trapped- state” (Chen *et al*., 1999; Sala-Rabanal *et al*., 2008) that is hypothesized to involve the binding competition between protons and substrates. In the mammalian transporters, hyperpolarization at acidic pH appears to have inactivating effects on *I*_PSS_ and steady-state properties, as in human and rat PepT2 the substrate-evoked current and substrate affinity decrease at very negative voltages at pH 5.5. In Atlantic salmon transporters, substrate affinity at hyperpolarizing potential decreases but it is not affected by pH; in the range of voltages tested the steady-state currents are not reduced by acidification and the *I*_PSS_ is recorded in the absence of the substrate and disappears in its presence. On the contrary, in the mammalian PepT2 the presence of external substrate seems to have a role in reducing the transporter inactivation by hyperpolarization because it increases the *I*_PSS_ and the maximal steady-state currents (*I*_max_) evoked by saturating substrate concentrations. The biophysical characterization of piscine PepT2 transporters and the data reported about the *I*_PSS_ can be the fundamental for future investigations on the relation between proton and substrate in the translocation steps of the peptide transporter family. In particular, it is of great interest that in all the PepT-type transporters functionally characterized so far, the residues identified by the recent structural model (Killer *et al*., 2021; Parker *et al*., 2021) as involved in the transporter activity, are conserved (**Fig. 1**). In the light of the results reported here, it will be important to investigate on residues that are not the main actors in transport activity but that could be responsible for the subtle changes in proton and substrate affinity, changes in substrate selectivity (Pieri *et al*., 2009; Xu *et al*., 2010; Margheritis *et al*., 2013) and to model their role in a specific transporter of interest.

## Conclusions

The Atlantic salmon represents the second teleost fish species (Ronnestad *et al*., 2010; Gomes *et al*., 2020) (this paper), after the zebrafish (Verri *et al*., 2003; Romano *et al*., 2006; Vacca *et al*., 2019), for which both PepT1- and PepT2-type transporters have been functionally characterized. Due to a salmonid specific WGD, two functional PepT2-type transporters operate in the Atlantic salmon which seems to have resulted in a split of functions with respect to the canonical situation described in mammals, in which a single PepT2 transporter operates in the whole organism. In this respect, our description of the Atlantic salmon, where one of the transporters (PepT2b) is expressed in kidney and mid-to-distal intestine, while the other (PepT2a) is expressed in the brain and gills, represents a novel paradigm in the vertebrates. Both transporters have specific transport traits, detailed here by biophysical and kinetic analysis. Moreover, the kinetic schemes previously proposed for two mammalian PepT2 transporters and the recently mechanistic structural models can qualitatively account for the transport structure and function. In a perspective, an effort is still needed to complete the local distribution of the two PepT2-type transporters in the various tissues/organs of the Atlantic salmon and demonstrate that these are expressed at the protein level. Also, the analysis of the regulation of these transporters remains open for future studies, as well as if and how they differentially respond to various external stimuli/environmental conditions, such as dietary nutrients, salinity, and temperature. Finally, the availability of detailed functional data from distant orthologs of the same protein certainly represents a key tool for investigating how single determinants can be responsible for subtle functional differences (Castagna *et al*., 2022), as the differences in sequence can easily be related to differences in the kinetic or biophysical parameters. From this point of view, the data here reported will surely be of help in the deep investigation on the structure-function relationships within the peptide transporters family.

## Author contributions

The experiments were performed in these laboratories: Laboratory of Cellular and Molecular Physiology, Department of Biotechnology and Life Sciences, University of Insubria, Varese (Italy); Department of Biological Sciences, University of Bergen, Bergen (Norway); Research Center for Aquaculture Systems, National Research Institute of Aquaculture, Japan Fisheries Research and Education Agency, Minami-ise, Mie (Japan); Laboratory of Applied Physiology, Department of Biological and Environmental Sciences and Technologies, University of Salento, Lecce (Italy).

FV, AG, KM, CR, RC, AB, TV and EB acquisition, analysis, and interpretation of data for the work; FV, EB, TV and IR design, drafting the work and revising it critically for important intellectual content. All authors have approved the final version of the manuscript; all authors agree to be accountable for all aspects of the work in ensuring that questions related to the accuracy or integrity of any part of the work are appropriately investigated and resolved; all persons designated as authors qualify for authorship, and all those who qualify for authorship are listed.

## Acknowledgments

The authors would like to thank Drs. Anne-Elise O. Jordal, Anders Aksnes, and Mali B. Hartviksen for technical assistance during sampling, Dr. Anders Aksnes (Cargill Innovation) for providing the fish, and Patrik Tang and Marius Takvam for providing the Atlantic salmon kidney and head-kidney RNA.

## Data availability statement

The Statistical Summary document contains data that support the findings of this study.

